# TRPML1 activation ameliorates lysosomal phenotypes in CLN3 deficient retinal pigment epithelial cells

**DOI:** 10.1101/2023.06.21.545896

**Authors:** D. Wünkhaus, R. Tang, K. Nyame, N. N. Laqtom, M. Schweizer, A. Scotto Rosato, E. K. Krogsæter, C. Wollnik, M. Abu-Remaileh, C. Grimm, G. Hermey, R. Kuhn, D. Gruber-Schoffnegger, S. Markmann

## Abstract

Mutations in the lysosomal membrane protein CLN3 cause Juvenile Neuronal Ceroid Lipofuscinosis (JNCL). Activation of the lysosomal ion channel TRPML1 has previously been shown to be beneficial in several neurodegenerative disease models. Here, we tested whether TRPML1 activation rescues disease-associated phenotypes in CLN3-deficient retinal pigment epithelial (ARPE-19 CLN3-KO) cells. ARPE-19 CLN3-KO cells accumulate LAMP1 positive organelles and show lysosomal storage of mitochondrial ATPase subunit C (SubC), globotriaosylceramide (Gb3), and glycerophosphodiesters (GPDs), whereas lysosomal bis(monoacylglycero)phosphate (BMP/LBPA) lipid levels were significantly decreased. Activation of TRPML1 reduced lysosomal storage of Gb3 and SubC but failed to restore BMP levels in CLN3-KO cells. TRPML1-mediated decrease of storage was TFEB-independent, and we identified TRPML1-mediated enhanced lysosomal exocytosis as a likely mechanism for clearing storage including GPDs. Therefore, ARPE-19 CLN3-KO cells represent a human cell model for CLN3 disease showing many of the described core lysosomal deficits, some of which can be improved using TRPML1 agonists.

## Introduction

Juvenile neuronal ceroid lipofuscinosis (JNCL), also named Batten or CLN3 disease, is the most common form of neuronal ceroid lipofuscinoses (NCLs), a group of rare inherited disorders that together represent the most prevalent neurodegenerative diseases in children^1, 2^. JNCL is caused by autosomal recessive mutations in the *CLN3* gene and manifests in blindness, seizures, progressive cognitive decline, and premature death of patients in their twenties to thirties^1, 3^. There is no disease-modifying therapy available for JNCL, and current treatment options only address symptoms.

Studies of CLN3 localization and function have been hampered by factors including cells expressing low levels of the *CLN3* gene, the hydrophobic nature of the protein, and the lack of animal models showing robust phenotypes seen in patients^3, 4^. Several studies suggest a role of CLN3 in endosomal-lysosomal compartments, autophagy, and in protein transport between the trans-Golgi network and lysosomes^5–8, 9, 10^. The CLN3 protein itself appears mainly localized in late endosomes and lysosomes. Earlier protein topology models predicted a six transmembrane domain lysosomal protein with both N- and C-terminus facing the cytoplasm^11, 12^. More recently, a protein structure with 11 transmembrane domains was proposed using Alphafold (https://alphafold.ebi.ac.uk/entry/Q13286) with the C-terminus facing the lysosomal lumen. CLN3 deficiency leads to lysosomal storage and accumulation of lipofuscin^5, 7, 13^, proteins such as mitochondrial ATPase subunit C (SubC)^13^, lipids including lyso-glycerophospholipids (lyso-GPL) and globotriaosylceramide (Gb3)^14^, and lipid breakdown products including glycerophosphodiesters (GPDs)^15^, while the levels of intralysosomal bis(monoacylglycerol)phosphates (BMPs) that are highly enriched in the membrane of intra-luminal vesicles (ILVs), are reduced^16^.

JNCL patients commonly first present with vision loss, indicating that retinal function is exquisitely sensitive to CLN3 dysfunction. Different studies imply functional defects also in the retinal pigment epithelium (RPE) of JNCL patients^17, 18^. Therefore, we generated and characterized a novel CLN3-deficient human ARPE-19 cell line (ARPE-19 CLN3-KO) that manifests many of the core including recently discovered novel CLN3 lysosomal deficits, such as accumulation of SubC, Gb3, and GPD, as well as reduced BMP levels. This cell line provides a novel and robust tool to investigate pharmacological agents and their potential to normalize CLN3-associated lysosomal and cellular core deficits.

Recent studies have demonstrated that activation of the endolysosomal Ca^2+^ channel TRPML1 can decrease accumulation of lysosomal storage material and relieve autophagic impairments in several lysosomal storage disorders (LSDs) and neurodegenerative diseases^19–24^. The endolysosomal TRPML1 channel is an important regulator of various Ca^2+^-dependent processes. These include endolysosomal trafficking, vesicle fusion/fission, the biogenesis, reformation and exocytosis of lysosomes, induction of autophagy, and lysosomal and autophagy-related gene expression controlled by the transcription factor EB (TFEB)^25–28^. Effects of TRPML1 activation on CLN3 disease-related phenotypes have not been evaluated so far.

To this end we used our novel cell model (ARPE-19 CLN3-KO) that recapitulates many pathological hallmarks of JNCL disease. We show that pharmacological activation of TRPML1 leads to a reduction of storage material without restoring BMP levels in CLN3-deficient lysosomes. Our data indicate an interplay of multiple mechanisms of TRPML1 action, including increased lysosomal exocytosis, nuclear TFEB translocation, and autophagy induction – all potentially contributing to the reduction of lysosomal storage burden and suggesting that TRPML1 agonists may have therapeutic potential in JNCL.

## Results

### CLN3 Batten disease cellular phenotypes in ARPE-19 CLN3-KO cells

Using CRISPR gene-editing, a novel CLN3 deficient ARPE-19 cell line (ARPE-19 CLN3-KO) was generated. This line carries an 8-nucleotide deletion in exon 3 of both CLN3 alleles. The deletion causes a shift in the reading frame with a premature stop codon at amino acid 156 (Suppl. Fig. S1 A-C). It also results in greatly reduced mRNA expression levels of the mutant transcript, likely due to nonsense-mediated mRNA decay (Suppl. Fig. S1 D).

In other cellular models of CLN3 disease, phenotypes characterized by storage material are most prominent in terminally differentiated, non-replicating cells^13, 29, 30^. Accordingly, to exacerbate phenotypes in ARPE-19 CLN3-KO cells, mutant and WT cells were treated with mitomycin C to induce cell cycle arrest (CCA)^22^. Unless explicitly stated, results described below will refer to ARPE-19 cells following CCA.

As compared to WT ARPE-19 cells, ARPE-19 CLN3-KO cells showed significant accumulation of two previously reported lysosomal storage products found in CLN3-deficient cells. These include the subunit C protein of the mitochondrial ATPase (SubC)^1, 13, 29^, and the lipid Globotriaosylceramide (Gb3). Gb3 is known to accumulate in Fabry’s disease^31^, and as recently shown^14, 30^, we confirmed that Gb3 accumulates in CLN3-KO RPE cells (Fig. 1 A-D).

**Figure 1:**
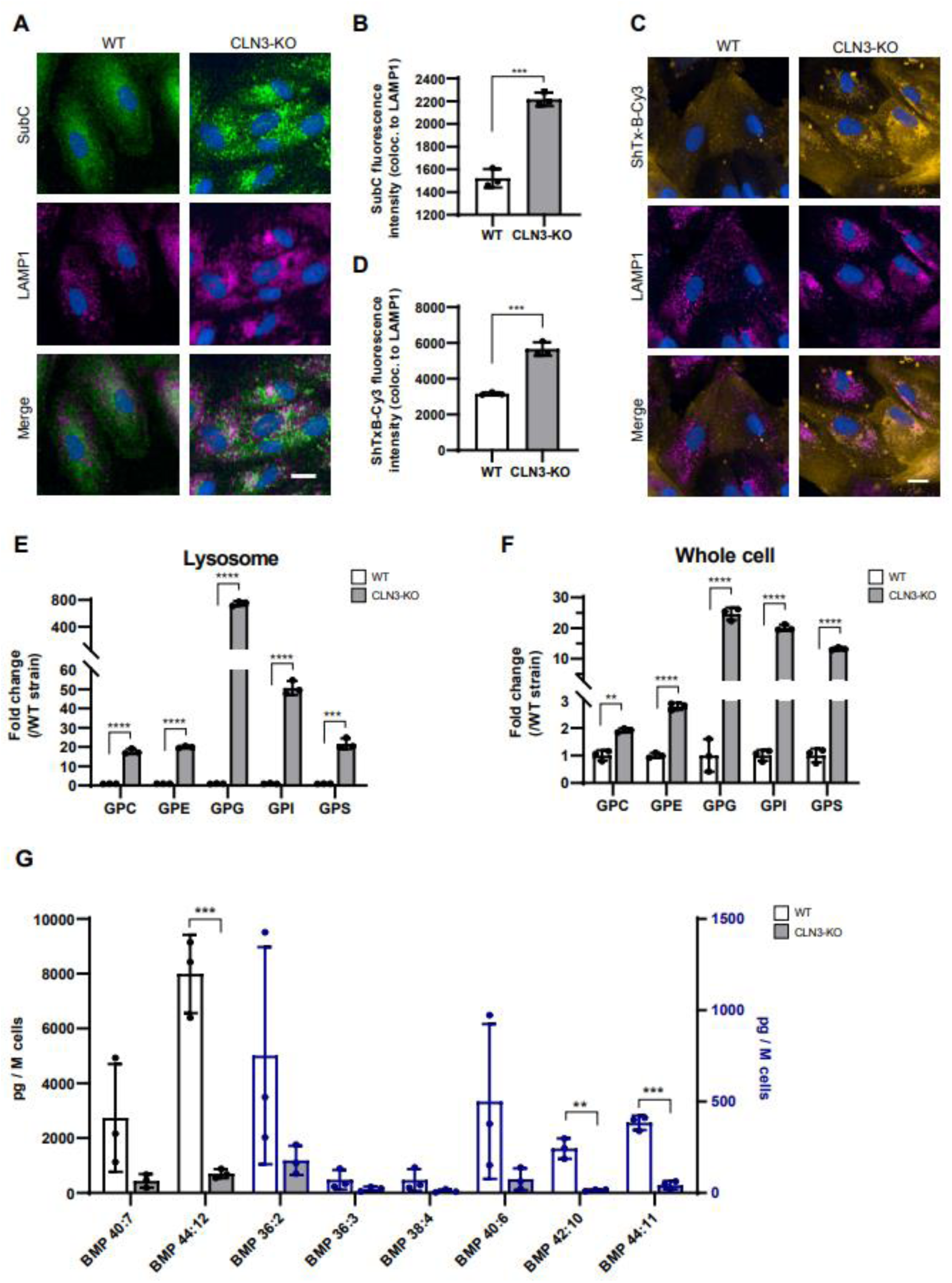
SubC, Gb3, GPDs and BMP/LBPA levels are altered in CLN3-KO cells. **A** Representative immunocytochemical images of SubC and LAMP1 in cell cycle arrested ARPE-19 WT and CLN3-KO cells. **B** Quantification of SubC fluorescence intensity colocalizing to LAMP1 (images shown in A). Representative plots from one experiment. In total three independent experiments were performed. Values are means ± SD. P-values were calculated by unpaired two-tailed Student’s T-test. **C** Representative immunocytochemical images of Gb3 labeling using ShTxB-Cy3 and LAMP1 in cell cycle arrested ARPE-19 WT and CLN3-KO cells. **D** Quantification of ShTxB-Cy3 fluorescence intensity colocalizing to LAMP1 (images shown in C) using data from one experiment. In total three independent experiments were performed. Values are means ± SD. P-values were calculated by unpaired two-tailed Student’s T-test. **E, F** Plots show the abundance of different glycerophosphodiesters: glycerophosphocholine (GPC), glycerophosphoethanolamine (GPE), glycerophosphoglycerol (GPG), glycerophosphoinositol (GPI), and glycerophosphoserine (GPS) in lysosomes purified using the lysoIP method (**E**) and in whole-cell lysates (**F**) of ARPE-19 WT and CLN3-KO cells with CCA. Plots summarize data from three independent experiments. P-values were calculated by individual unpaired two-tailed student’s t-test for each GPD. **G** Plot shows levels of different BMP species, detected in cell lysates of cell cycle arrested ARPE-19 WT and CLN3-KO cells using mass spectrometry. Values are means ± SD of three independent experiments. P-values were calculated by individual unpaired two-tailed Student’s t-test for each BMP species.

Using lysosome immunoprecipitation (lysoIP), glycerophosphodiesters (GPDs) were recently identified as a group of glycerophospholipid metabolites that selectively accumulate inside lysosomes of CLN3-deficient human and mouse cells^15^. In agreement, ARPE-19 CLN3-KO cells showed massive accumulation of GPDs including GPC, GPE, GPG, GPI and GPS in lysosomes (Fig. 1 E). Interestingly, mass spectrometry analysis of whole cell extracts also showed significantly elevated GPD levels (Fig. 1F), suggesting that the changes in lysosomal levels are readily detected in whole cell measurements.

Intralysosomal vesicle (ILV) membranes are enriched in a typical anionic lipid family known as BMPs. BMPs account for 15-20 Mol% of total phospholipids in the late endosomal/lysosomal compartment, and are not detected in other cellular compartments^32, 33^. BMPs located in the internal membranes of lysosomes interact with activator proteins and are necessary to facilitate lipid degradation^34, 35^. BMP levels are reduced in some LSDs and corresponding animal models including CLN3^15, 16^ and progranulin (GRN/CLN11)-deficiency^36^. Increasing BMP levels has been shown to ameliorate storage phenotypes such as cholesterol overload in NPC1-/- cells^37, 38^ and gangliosidosis in GRN-deficient cells^36^. In agreement with previous findings^15, 16^, ARPE-19 CLN3-KO cells as compared to their isogenic WT counterparts showed a significant depletion of BMP lipid species (Fig. 1 G).

Like observations in other CLN3-deficient cell types, ARPE-19 CLN3-KO cells showed increased levels of the lysosomal membrane proteins LAMP1, LAMP2 and NPC1, as well as acidic compartment staining by LysoTracker, in cells before as well as after CCA (Suppl. Fig. S2 A-E). Levels of the corresponding mRNAs (LAMP1, LAMP2, NPC1) were not significantly changed in CLN3-KO versus WT cells (Suppl. Fig. S2 F-H). Electron microscopy (EM) analysis corroborated the LAMP1 immunofluorescence results. EM analysis revealed significant increases in lysosomal size, number, and total lysosomal compartment area per cell in CLN3-KO as compared to WT cells (Fig. 2 A, B). The morphological appearance of CLN3-deficient lysosomes was heterogenous including both empty vacuoles as well as lysosomes containing small internal electron-dense structures or ‘fingerprint-like’ inclusions as described in JNCL^13, 29^.

**Figure 2:**
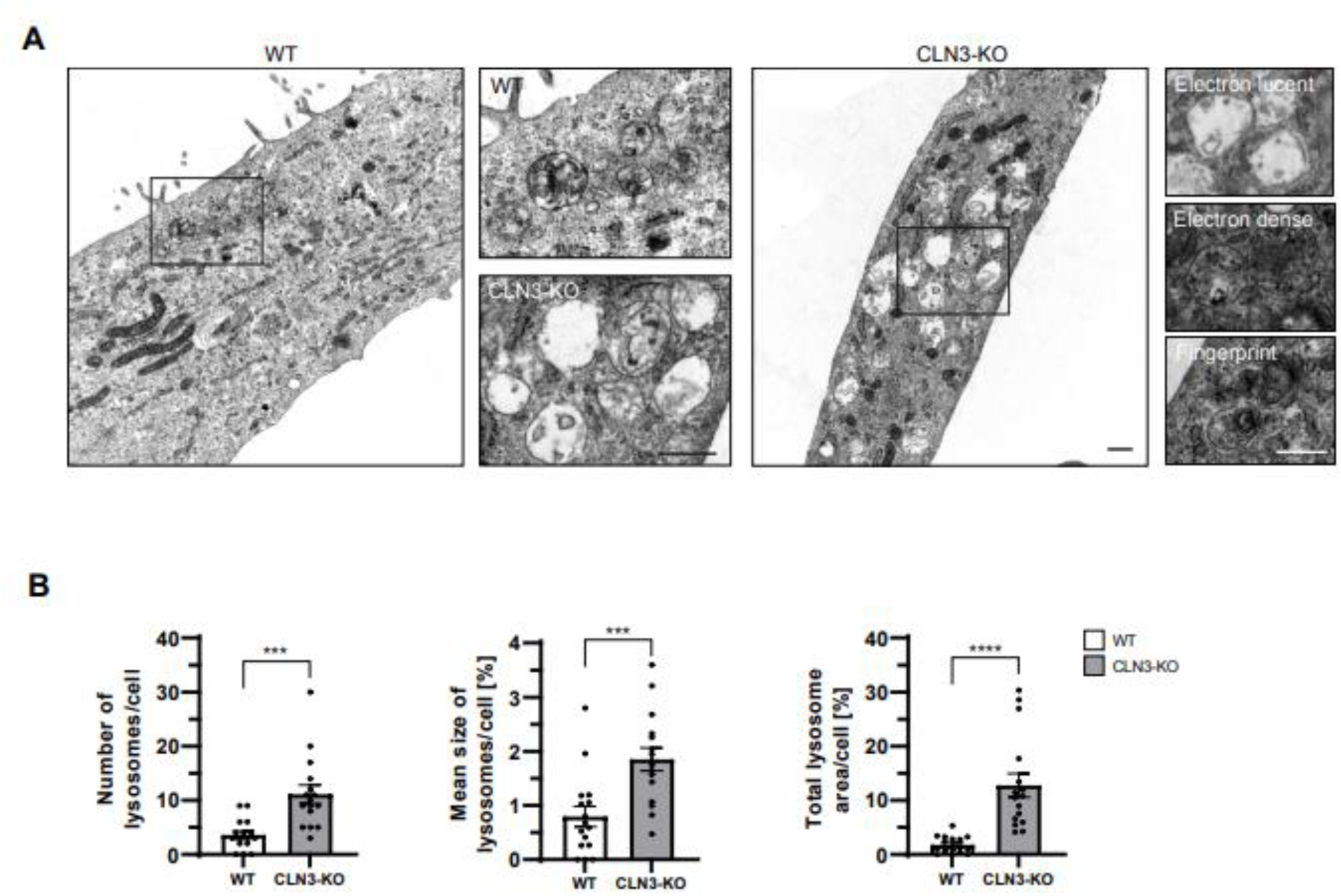
Increased lysosomal number, size and area in CLN3-KO cells. **A** Representative electron microscopy images of cell cycle arrested ARPE-19 WT and CLN3-KO cells. **B** Plots show quantification of lysosomal number, size and total lysosomal area per cell from 16 electron microscopy images. Values are means ± SEM. Scale bar: 500 nm. P-values were calculated by unpaired two-tailed Student’s T-test.

### TRPML1 activation reduces lysosomal storage in ARPE-19 CLN3-KO cells

To ensure that ARPE-19 CLN3-KO cells express and respond to TRPML1 agonists, we determined TRPML1 mRNA levels and monitored agonist-evoked TRPML1 ion channel currents in CLN3-KO cells. TRPML1 mRNA levels were similar between untreated WT and CLN3-KO cells. Likewise, after stimulation with the agonist ML-SA5^22^ (Fig. 3 A). Lysosomal patch clamp analysis using the TRPML1 agonist ML-SA1^36^ revealed the presence of TRPML1 currents in lysosomes of both WT and ARPE-19 CLN3-KO cells (Fig. 3 B-F). Agonist-evoked TRPML1 currents were fully blocked by treatment with the TRPML1 antagonist ML-SI3^39, 40^ (Fig. 3 B-F).

**Figure 3:**
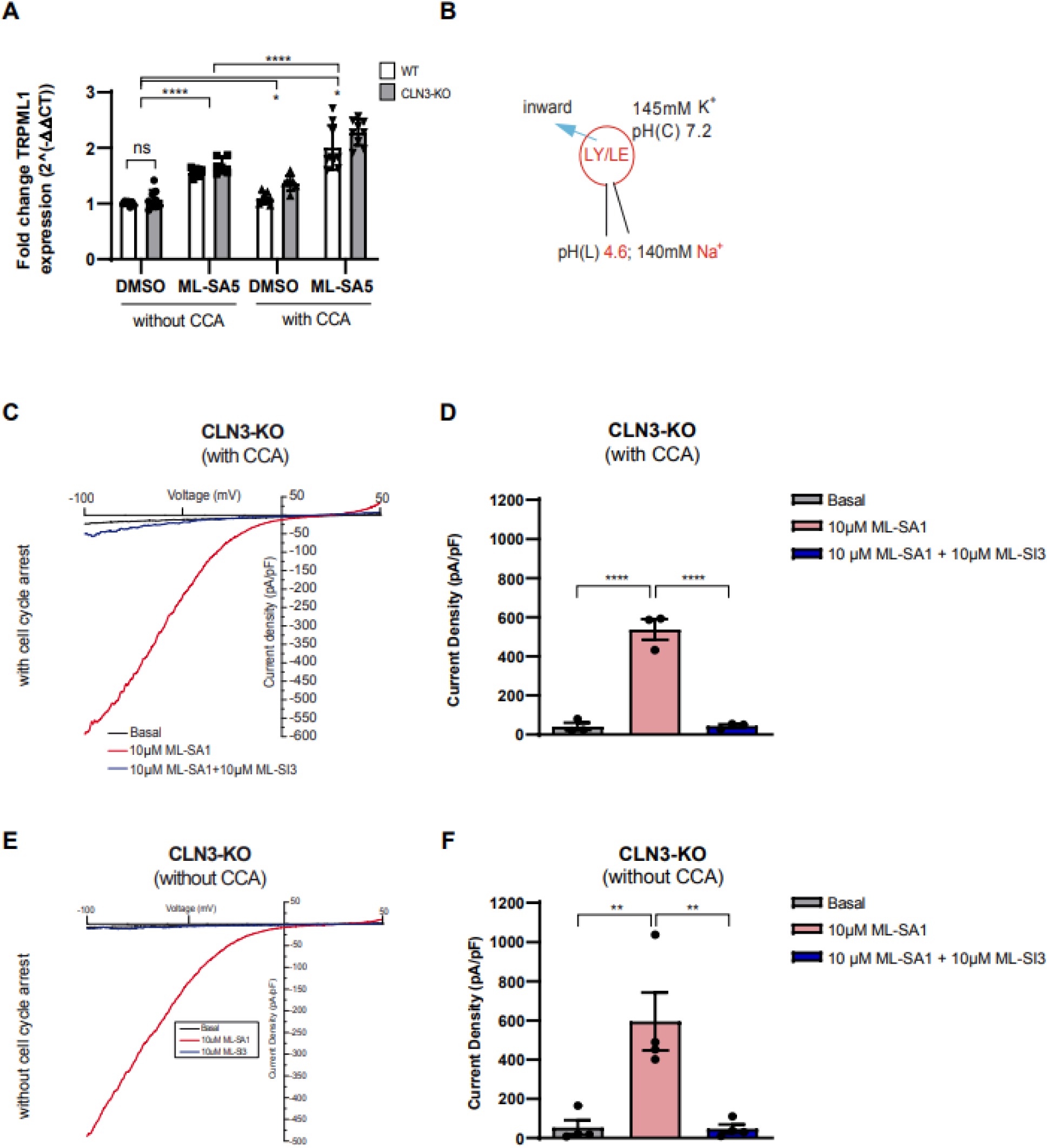
TRPML1 expression and currents in WT and CLN3-KO cells. **A** Relative mRNA expression levels of TRPML1, normalized to Peptidyl-prolyl cis-trans isomerase B (PPIB) and WT DMSO treated cells without CCA. Cells were treated for 24 h with 30 µM ML-SA5, with and without cell cycle arrest (CCA). Values are means ± SD, n=3 independent experiments. P-values calculated by two-way ANOVA coupled with multiple comparisons using Tukey test. **B** Scheme of lysosomal patch clamp technique, showing main buffer components. Note the pH difference between the lysosome pH(L)=4.6 and the cellular surrounding pH(C)=7.2. **C** Representative TRPML1 currents with agonist ML-SA1 and antagonist ML-SI3 treatment in cell cycle arrested ARPE-19 CLN3-KO cells, measured with whole endolysosomal patch clamp technique. **D** Plot shows current densities of three independently patched lysosomes from ARPE-19 CLN3-KO cells under CCA. P-values were calculated by one-way ANOVA coupled with multiple comparisons using Tukey test. **E** Representative TRPML1 currents with agonist ML-SA1 and antagonist ML-SI3 treatment of non-CCA ARPE-19 CLN3-KO cells, measured with whole endolysosomal patch clamp technique. **F** Plot shows current densities of four independently patched lysosomes from ARPE-19 CLN3-KO cells without cell cycle arrest. P-values were calculated by one-way ANOVA coupled with multiple comparisons using Tukey test. ns, nonsignificant<0.1234, *p-value<0.0332; **p<0.0021, ***p-value<0.0002, ****p-value<0.0001.

To test effects of TRPML1 agonist treatment on CLN3 deficiency-associated lysosomal storage phenotypes in ARPE-19 cells, we used instead of ML-SA1 the more potent TRPML1 agonist ML-SA5^22^.

Treatment of CLN3-KO cells with ML-SA5 significantly decreased SubC accumulation in a concentration-dependent fashion. The decrease was already detectable at 6 hours after drug administration but more pronounced after 48 hours (Fig. 4 A-C, Suppl. Fig. S3 A). Gb3 accumulation, visualized using Cy3-labeled Shiga toxin (ShTx-B-Cy3), was also decreased in a ML-SA5 concentration-dependent manner (Fig. 4 D-E). Analysis using electron microscopy of CLN3-KO cells treated with ML-SA5 showed a significant reduction in the total surface area of lysosomes, but not their numbers (Fig 4 F-G, Suppl. Fig. S3 B-C). ML-SA5 increased LAMP1 mRNA levels to the same extent in both genotypes (Suppl. Fig. S3 D), while LAMP1 protein levels remained largely unchanged (Suppl. Fig. S3 E-F).

**Figure 4:**
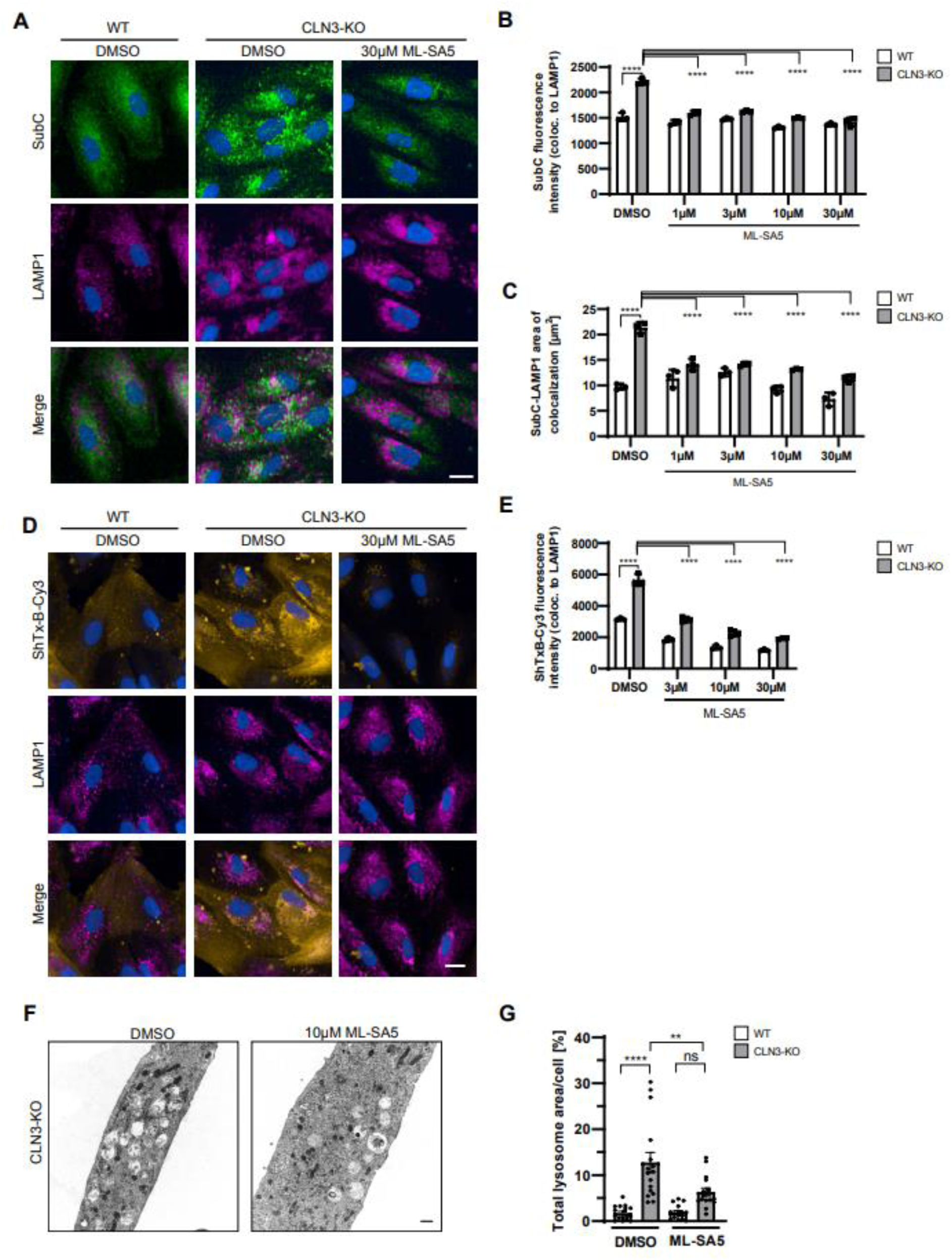
TRPML1 activation ameliorates phenotypes of CLN3-KO cells. **A** Representative immunocytochemical images of SubC (green) and LAMP1 (magenta) in cell cycle arrested ARPE-19 WT and CLN3-KO cells. ML-SA5 was added for 48 h, 7 days after induction of CCA. Scale bar: 20 µm. **B, C** Quantification of immunocytochemical images shown in A (**B)** for SubC fluorescence intensity and (**C**) area of SubC-LAMP1 colocalization. Representative plots from one experiment. In total three independent experiments were performed. Values are means ± SD. **D** Representative immunocytochemical images of Gb3 labeling using ShTxB-Cy3 (yellow) and LAMP1 (magenta) in cell cycle arrested ARPE-19 WT and CLN3-KO cells. Cells were treated with ML-SA5 for 48 h, 7 days after induction of CCA. Scale bar: 20 µm. **E** Quantification of ShTxB-Cy3 fluorescence intensity (images shown in D) using data from one experiment. In total three independent experiments were performed. Values are means ± SD. **F** Representative electron microscopy images of ARPE-19 CLN3-KO cells under cell cycle arrest. Cells were treated with 10 µM ML-SA5 for 48 h, 7 days after induction of CCA. Scale bar: 500 nm. **G** Plots show quantification of total lysosome area per cell from 16 electron microscopy images. Values are means ± SEM. P-values calculated by two-way ANOVA coupled with multiple comparisons using Tukey test.

Treatment with ML-SA5 also resulted in a significant, albeit small reduction of all the GPD species that accumulate inside the lysosomes of CLN3-KO cells. ML-SA5 was not able to normalize GPD levels back to WT levels (Fig. 5). Changes in GPD levels were measured at two time points (90 min and 72 hours) after treatment with ML-SA5. We observed that most GPDs were already significantly reduced after 90 min. Some GPDs were decreased further by 72 hours. The same trend was detected in whole-cell lysates with overall lower GPD levels and only GPI showing a further decrease at 72 hours as compared to 90 min (Suppl. Fig. S4).

**Figure 5:**
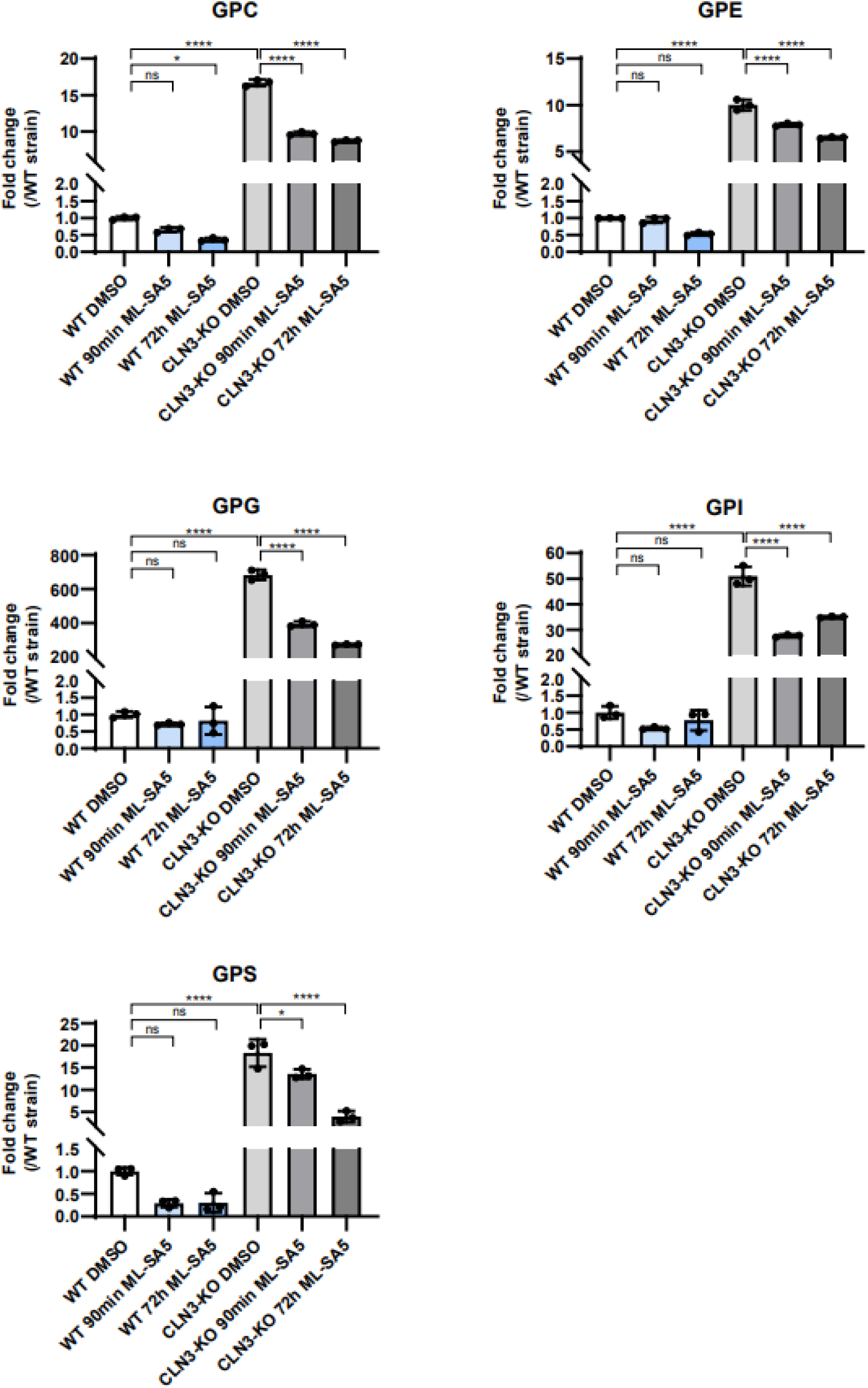
Effect of TRPML1 activation on lysosomal GPD levels in WT and CLN3-KO cells. Plots show the fold change of the different GPD species in isolated lysosomes from ARPE-19 WT and CLN3-KO cells expressing the lysosomal tag 3xHA-TMEM192. Cells were under CCA and treated with 10 µM ML-SA5 for 90min and 72h. Values are means ± SD of three independent experiments. P-values were calculated by one-way ANOVA coupled with multiple comparisons using Tukey test.

ML-SA5 seemed to slightly although non-significantly increase BMP levels in both genotypes but the drug failed to normalize BMP levels in CLN3-KO cells back to the levels seen in WT cells (Fig. 6). Likewise in cells without CCA (Suppl. Fig. S5). TRPML1-induced changes in BMP levels therefore unlikely provide a mechanism and an explanation for the beneficial effects of ML-SA5 observed on storage phenotypes in CLN3-KO cells including SubC, Gb3 and different GPDs.

**Figure 6:**
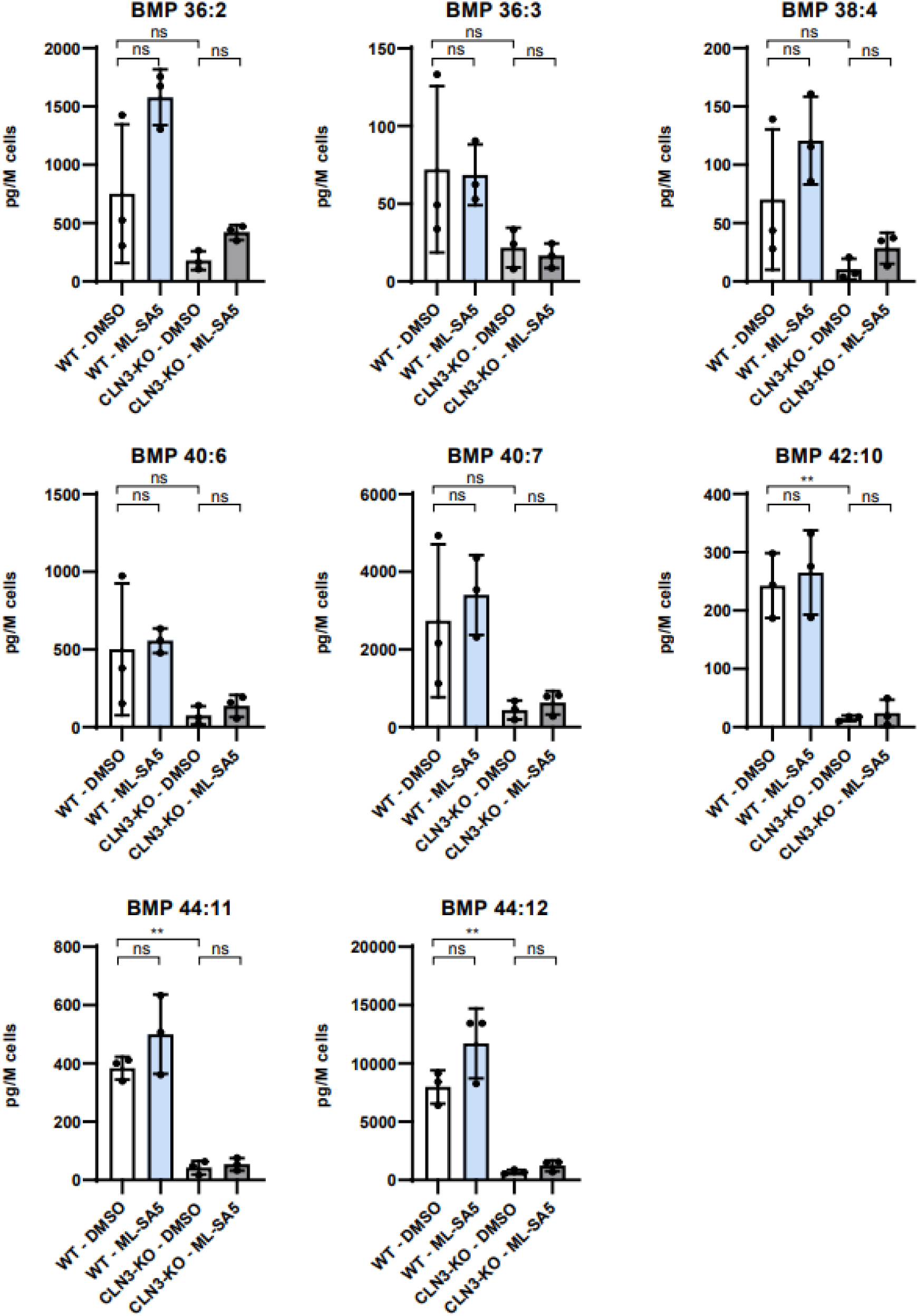
Effect of TRPML1 activation on BMP levels in WT and CLN3-KO cells. Plots show the abundance of BMP species in cell lysates of ARPE-19 WT and CLN3-KO cells with CCA and treatment with 3 µM ML-SA5 for 48h. Values are means ± SD of three independent experiments. P-values were calculated by one-way ANOVA coupled with multiple comparisons using Tukey test.

### TRPML1 activation induces TFEB translocation, but SubC and Gb3 reduction are TFEB-independent

TRPML1 has been described to regulate endolysosomal vesicle homeostasis, TFEB-dependent lysosomal biogenesis, enhance autophagy, CaMKKβ/VPS34-dependent and TFEB-independent induction of phagophore formation, lysosomal exocytosis, as well as vesicle fusion. Most of these mechanisms require its function as a lysosomal Ca^2+^ release channel, and some but not all require also TFEB^25–28^. TRPML1-mediated lysosomal Ca^2+^ release **triggers** dephosphorylation and nuclear translocation of the transcription factor EB (TFEB)^28^. TFEB is a master regulator of lysosomal biogenesis, controlling expression of lysosomal and autophagy-related genes containing the CLEAR (Coordinated Lysosomal Expression and Regulation) element^41^. ML-SA5 induced TFEB nuclear translocation to the same extent in both WT and CLN3-KO cells. This is reflected both by increased nuclear translocation by immunocytochemistry, and by the downward molecular weight shift of the dephosphorylated protein in western blot analysis (Fig. 7 A, B, C). For reference, the mTORC1 inhibitor Torin1 stimulates TRPML1-independent TFEB nuclear translocation and showed comparable effects (Fig. 7 A, B, C). TFEB protein and mRNA levels were not altered in CLN3-KO relative to WT cells (Fig. 7 C, D) but note that CCA increased TFEB mRNA levels to a similar extent in both WT and CLN3-KO cells (Fig S7). Of note, ML-SA5 and Torin1 also induced TFEB nuclear translocation in ARPE-19 cells without CCA, but to a smaller extent than in cells after CCA (Suppl. Fig. S6 A, B). Finally, mRNA levels of several TFEB regulated lysosomal and autophagy related genes were analyzed. ML-SA5 treatment caused significant increases in several of these TFEB-responsive genes in both WT and CLN3-KO cells with and without CCA (Fig. S7). These data confirm that TRPML1 agonist-induced TFEB-mediated gene expression changes in the autophagy-lysosomal pathway are not compromised by lack of CLN3.

**Figure 7:**
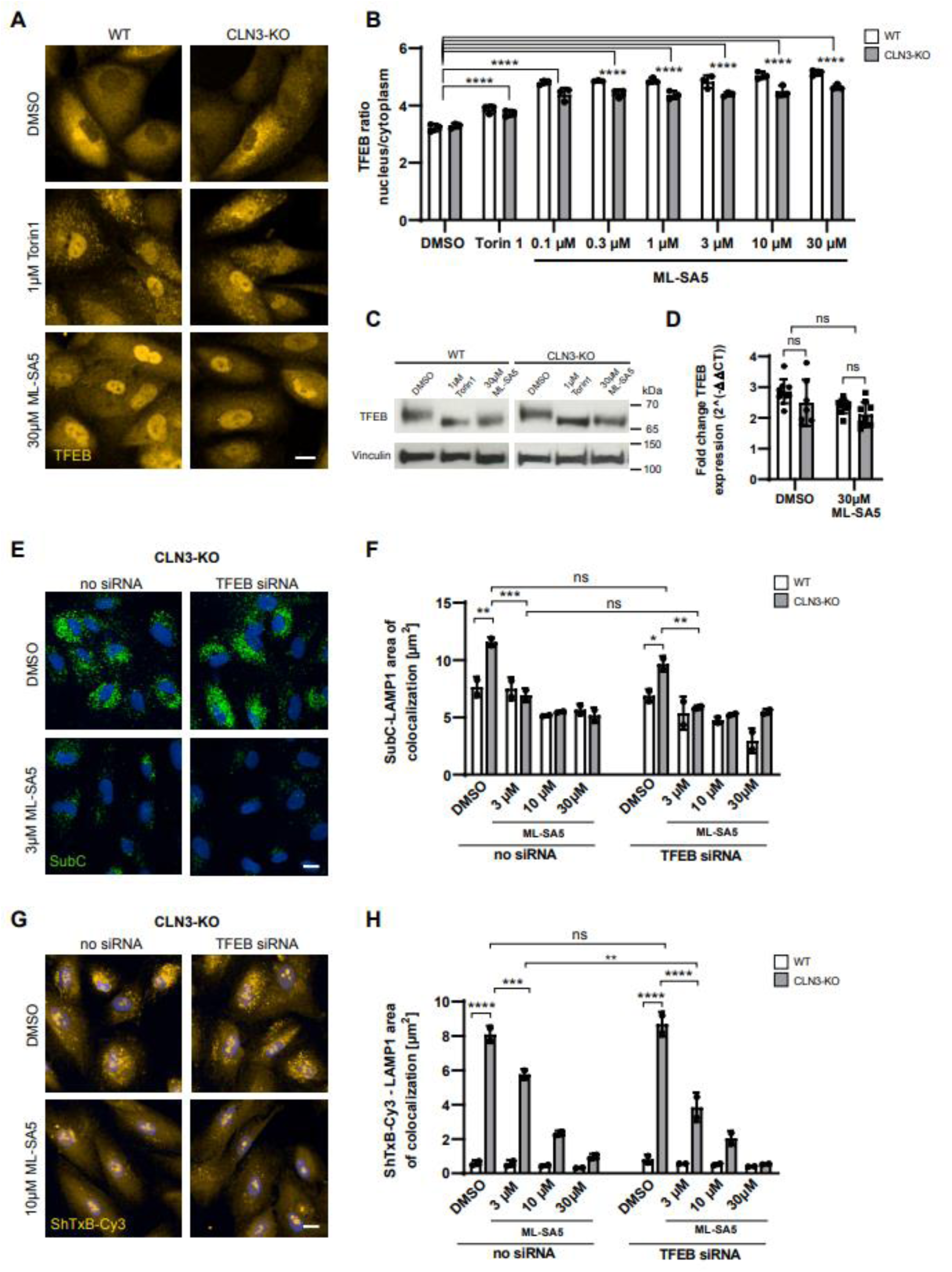
Effect of TRPML1 activation on TFEB expression and nuclear translocation and TFEB knockdown in WT and CLN3-KO cells. **A** Representative immunocytochemical images showing TFEB nuclear translocation after treatment of cell cycle arrested ARPE-19 WT and CLN3-KO cells for 2 h with 1 µM Torin1 and 30 µM ML-SA5. Scale bar: 20 µm. **B** Quantification of images shown in A. Plot shows TFEB nuclear/cytosol ratio of one representative experiment. In total three independent experiments were performed. Values are means ± SD. **C** Immunoblot analysis of total cellular TFEB levels. Cell cycle arrested ARPE-19 WT and CLN3-KO were treated with mTORC-inhibitor Torin1 and TRPML1 agonist ML-SA5 for 2 h. Treatment leads to TFEB dephosphorylation and nuclear translocation, which is reflected by a size shift. **D** Relative mRNA expression levels of TFEB, normalized to PPIB and WT DMSO treated cells without CCA. Cells were treated for 24 h with 30 µM ML-SA5. Values shown are means ± SD, n=3 independent experiments. **E, G** Representative immunocytochemical images of SubC (**E**) and Shigatoxin labeled Gb3 (**G**) after siRNA mediated knockdown of TFEB in CLN3-KO cells. Scale bar: 20 µm. **F, H** Quantification of images shown in E and G. Plots show SubC-LAMP1 (**E**) and ShTxB-Cy3-LAMP1 (**G**) area of colocalization of cell cycle arrested ARPE-19 cells with and without addition of TFEB siRNAs, followed by ML-SA5 treatment for 48 h. Representative plots from one experiment. In total three independent experiments were performed. Values are means ± SD. P-values were calculated by a two-way ANOVA coupled with multiple comparisons using Tukey test.

To determine whether the TRPML1-mediated reduction of SubC and Gb3 accumulation is dependent on TFEB, siRNA-mediated TFEB knock-down was performed (Suppl. Fig. S6 C). We observed comparable decreases of SubC and Gb3 accumulation after treatment with ML-SA5 in CCA ARPE-19 CLN3-KO cells with and without TFEB knock-down (Fig. 7 E-H). This indicates that TRPML1-mediated reduction of SubC and Gb3 storage is TFEB-independent.

### TRPML1 activation induces autophagy in ARPE-19 WT and CLN3-KO cells

Several studies have documented autophagic defects in CLN3-deficient cells^6, 9, 29, 42^. TRPML1 activation induces both phagophore formation through activation of CaMKKβ/VPS34^27^, as well as TFEB-mediated upregulation of CLEAR genes and lysosomal biogenesis^43^. To determine whether TRPML1-mediated effects on autophagosome and autolysosome formation are different in WT and CLN3-KO cells we monitored LC3-II levels. LC3-II is the lipidated form of LC3 and generated by the conjugation of cytosolic LC3-I to phosphatidylethanolamine (PE). LC3-II is recruited to the surface of nascent autophagosomes, and is therefore commonly used as a marker for autophagosomes^44^. 30 min after treatment with ML-SA5, both WT and CLN3-deficient cells showed a rapid increase in the levels of LC3-II protein (Fig. 8 A-B), in agreement with previously reported results^27^. LC3 mRNA levels were similar and unchanged in CLN3-KO and WT cells (Fig. 8 C). Immunofluorescence analysis confirmed a rapid increase of LC3 punctate structures (Fig 8 E-D). Treatment with ML-SA5 also significantly increased the number of LC3 structures colocalizing with LAMP1 representing autolysosomes (Fig. 8 D-F). Cells without CCA showed similar results (Suppl. Fig. S8 A-F). Taken together, these data show that TRPML1-mediated effects on autophagosome and autolysosome formation are similar in WT and CLN3-KO cells and therefore not significantly perturbed by CLN3 deficiency.

**Figure 8:**
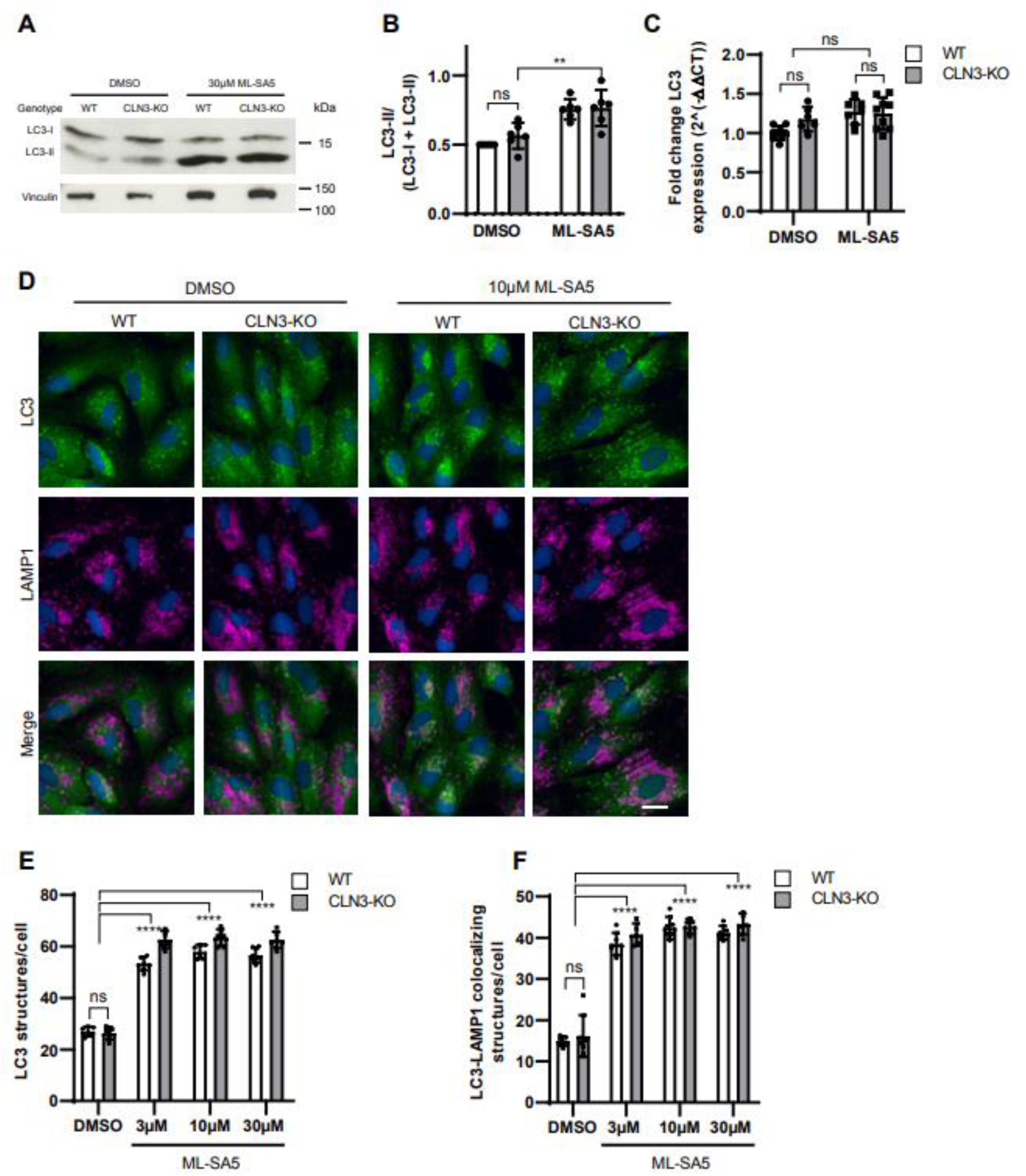
Effect of TRPML1 activation on autophagy in WT and CLN3-KO cells. **A** Immunoblot analysis of LC3. Cell cycle arrested ARPE-19 WT and CLN3-KO cells were treated for 30 min with 30 µM ML-SA5. **B** Quantification of LC3-II/(LC3-I + LC3-II) levels normalized to Vinculin, n=6 independent experiments. **C** Relative mRNA expression of LC3 normalized to Peptidyl-prolyl cis-trans isomerase B (PPIB) and DMSO treated WT cells without CCA. Cell cycle arrested ARPE-19 cells were treated for 24 h with 30 µM ML-SA5. Values are means ± SD, n=3 independent experiments. **D** Representative immunocytochemical images of LC3 and LAMP1 localization in cell cycle arrested ARPE-19 WT and CLN3-KO cells. Scale bar: 20 µm. **E, F** Plots represent the number of LC3 structures/cell (**E**) and LC3-LAMP1 colocalizing structures/per cell (**F**). Cells were treated with ML-SA5 for 30 min (**D-F**). Plots summarize data from one experiment. In total 3 independent experiments were performed. Values are means ± SD.

### TRPML1 activation induces lysosomal exocytosis in WT and CLN3-KO cells

Next, we investigated whether lysosomal exocytosis might be a mechanism by which TRPML1 agonists can clear GPDs and thereby reduce lysosomal storage in CLN3-deficient cells. During lysosomal exocytosis, lysosomal membranes fuse with the plasma membrane. This enables plasma membrane repair, a process proposed to underly therapeutic effects of ML-SA5 in a DMD mouse model of muscle disease^22^, as well as clearance of lysosomal content into the extracellular space. The fusion of lysosomal membranes with the plasma membrane is Ca^2+^-dependent and triggered by TRPML1-mediated lysosomal Ca^2+^ release^45^. To probe TRPML1-induced lysosomal exocytosis in the ARPE-19 cells, we used FACS to monitor the expression of a LAMP1 luminal epitope on the cell surface. As a positive control, cells were treated with the ionophore ionomycin, which triggers a massive increase of cytoplasmic Ca^2+^ levels and induces TRPML1-independent lysosomal exocytosis^46^. A 90 min treatment with ML-SA5 induced lysosomal exocytosis in a concentration-dependent fashion in both WT and CLN3-KO cells. 20-40% of all cells showed cell surface expression of the luminal LAMP1 epitope (Fig. 9 A-E).

**Figure 9:**
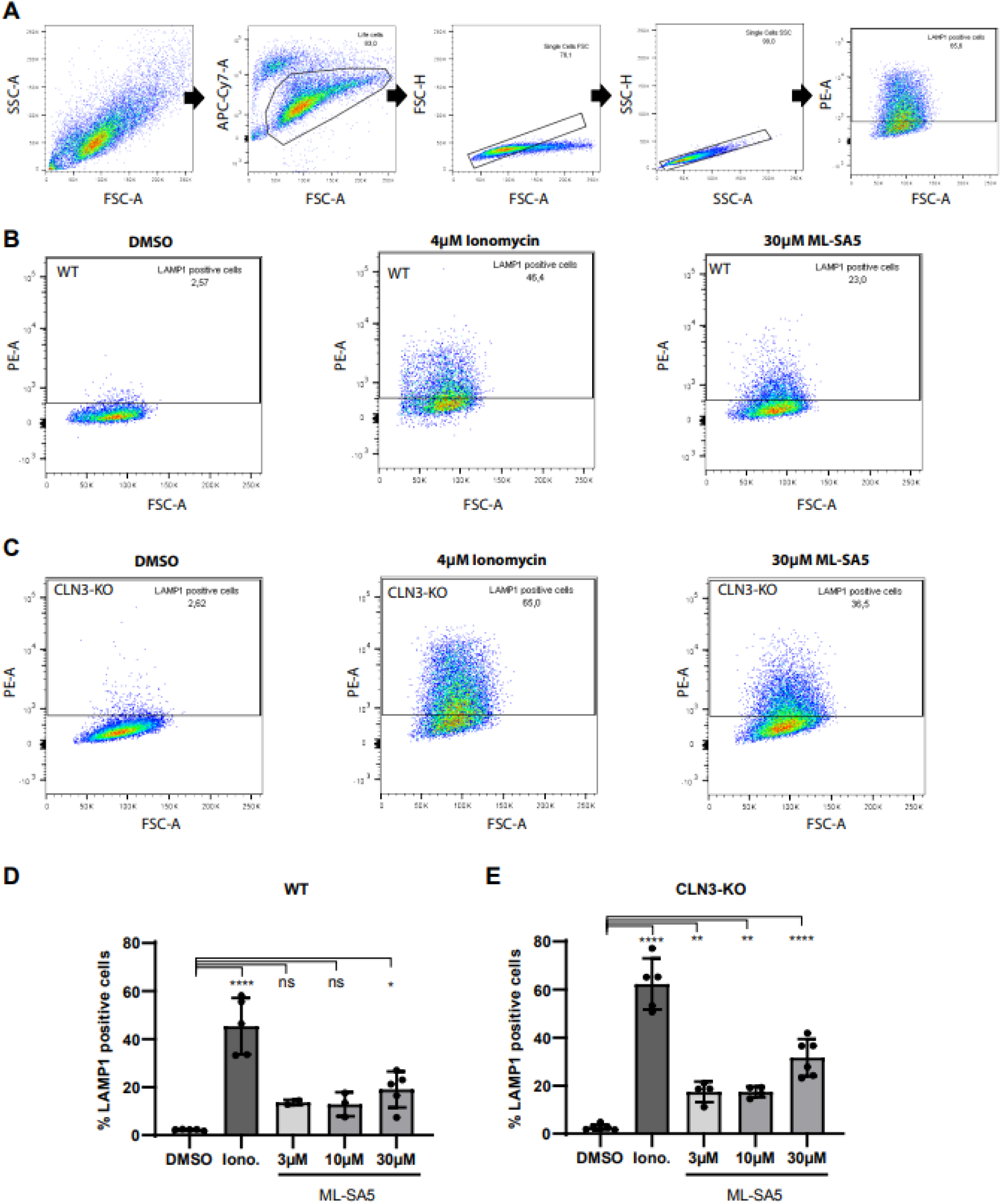
Effect of TRPML1 activation on lysosomal exocytosis in WT and CLN3-KO cells. **A** Gating strategy used to identify LAMP1 positive cells by flow cytometry. **B, C** Exemplary flow cytometry plots in cell cycle arrested ARPE-19 WT (**B**) and CLN3-KO (**C**) cells. **D, E** Plots represent the percentage of cell cycle arrested ARPE-19 WT (**D**) and CLN3-KO (**E**) cells showing plasma membrane LAMP1 staining. For reference, cells were treated for 15 min with Ionomycin (Iono.). ML-SA5 treatment was performed for 90 min. Shown are individual values from single experiments ± SD of all experiments, n ranges from 3-6 independent experiments.

The observed TRPML1-mediated lysosomal exocytosis in ARPE-19 WT and CLN3-KO cells is in accordance with the reduction in GPDs observed 90 min after ML-SA5 treatment (Fig. 5), suggesting that lysosomal exocytosis might be one of the mechanisms by which TRPML1 agonists clear lysosomal storage in CLN3-deficient cells.

## Discussion

In this study, we demonstrated that the activation of the endolysosomal cation channel TRPML1 rescues several lysosomal storage phenotypes in CLN3 deficient ARPE-19 cells, a retinal epithelium JNCL cell model. CLN3-deficiency is the cause of JNCL, a rare form of childhood dementia. Rapid and progressive retinal degeneration is the most consistent earliest clinical symptom in JNCL patients and diagnosis is often delayed, because other Batten disease clinical symptoms show significant inter-individual variability and manifest later^47^. No disease-modifying treatments currently exist and biomarkers are urgently needed to facilitate clinical trials^48, 49^. Robust cell pathological phenotypes in disease-relevant cell models and a better understanding of which treatments normalize CLN3 hallmarks may help to pave the road for a better translation of treatments and biomarkers from preclinic to clinic.

As compared to their isogenic WT counterparts, ARPE-19 CLN3-KO cells show lysosomal accumulation of SubC protein, and the lipid Globotriaosylceramide (Gb3). Gb3 accumulates in Fabry’s disease (alpha-galactosidase A deficiency)^31^, and was recently shown to accumulate also in another independently generated CLN3-KO as well as a CLN7 RPE cell line, as well as in brain neurons in vivo^14, 30^. In addition, ARPE-19 CLN3-KO cells showed increased levels of the lysosomal membrane proteins LAMP1, LAMP2 and NPC1 without changes in the levels of their corresponding mRNAs. Electron microscopy (EM) analysis revealed significant increases in lysosomal size, number, and total lysosomal compartment area per cell in CLN3-KO as compared to WT cells. CLN3-KO lysosomes appeared either empty or contained small electron-dense structures or ‘fingerprint-like’ inclusions as described in other JNCL cell types^13, 29^.

Importantly, our novel ARPE-19 CLN3-KO cell model showed massive accumulation of glycerophosphodiesters (GPDs) including GPC, GPE, GPG, GPI and GPS. GPDs were recently identified as degradation products of glycerophospholipids that accumulate inside lysosomes of CLN3-deficient human and mouse cells^15^. Interestingly, ARPE-19 CLN3-KO cells also showed significantly increased GPD levels in whole cell extracts, a feature that may greatly facilitate the use of these cells in screening efforts looking for normalizing omics changes and storage phenotypes.

In agreement with recent findings in CLN3-deficient human cell models and mice^15, 16^, we confirmed that BMP levels are greatly reduced in ARPE-19 CLN3-KO cells. These anionic lipids are highly enriched in intra-lysosomal vesicle membranes, where BMPs interact with activator proteins and enzymes to facilitate lipid degradation^34, 35^. BMP levels are reduced in some LSDs and corresponding animal models including CLN3-deficiency^15, 16^ and progranulin (GRN/CLN11)-deficiency^36^. Increasing BMP levels has been shown to ameliorate storage phenotypes such as cholesterol overload in NPC1^-/-^ cells^37, 38^, and gangliosidosis in GRN-deficient cells^36^. Reduced levels of BMPs in CLN3-KO lysosomes may therefore contribute to the lysosomal storage phenotypes observed in these cells.

After confirming that TRPML1 is expressed and functional in ARP19 CLN3-KO cells, we showed that treatment with ML-SA5 concentration-dependently decreased SubC and Gb3 accumulation, and significantly reduced intra-lysosomal GPD levels. The treatment did not rescue GPD levels to the low levels seen in WT cells. Reduction in GPD levels were observed after 90 min treatment, and for several GPDs the levels continued to decrease further by 72 hours. Finally, ML-SA5 had no significant effect on BMP levels, which remained low in CLN3-KO cells. Therefore, ML-SA5-mediated effects on reducing storage seem unlikely caused via a mechanism involving BMP regulation.

Several, but not all downstream effects of TRPML1 activation and lysosomal Ca^2+^ release involve the transcription factor TFEB. These include enhancing lysosomal biogenesis, induction of CLEAR genes and autophagy, and facilitating lysosomal exocytosis. TRPML1-mediated Ca^2+^ release triggers the dephosphorylation of TFEB and its nuclear translocation. We showed that ML-SA5 triggers TFEB nuclear translocation to the same extent in WT and ARPE-19 CLN3-KO cells, and also results in similar increases of mRNA expression of several TFEB-regulated lysosomal and autophagy-related genes. Furthermore, TRPML1-mediated effects on autophagosome and autolysosome formation were of similar magnitude in WT and CLN3-KO cells. Therefore, the lack of CLN3 does not seem to compromise TFEB-mediated changes in response to TRPML1 activity, nor does it significantly affect the process of autophagosome and autolysosome formation. To determine, whether TRPML1-mediated reduction of storage phenotypes like SubC and Gb3 accumulation requires TFEB, we performed siRNA-mediated knockdown in CCA ARPE-19 CLN3-KO cells. TFEB knockdown failed to prevent rescue of storage mediated by TRPML1 activation, suggesting that the underlying mechanism of storage clearance does not require TFEB.

The rapid lowering of intra-lysosomal GPD levels after treating CCA ARPE-19 CLN3-KO for 90 min with ML-SA5 suggests that lysosomal exocytosis might be a key mechanism by which TRPML1 activation reduces storage burden. To measure lysosomal exocytosis, we monitored the expression of a LAMP1 luminal epitope on the cell surface by FACS. A 90 min treatment with ML-SA5 triggered a robust expression of LAMP1 on the cell surface, in line with our findings of reducing GPD levels at this time point by a mechanism involving exocytosis.

Altogether, our work adds new insights into the cellular implications of CLN3 mutations in retinal pigment epithelial cells. Based on the reduction of diverse lysosomal accumulation materials and the investigation of several Ca^2+^-dependent mechanisms, we provide strong evidence, that activation of the ion channel TRPML1 can ameliorate the phenotype of CLN3 defective cells. Future work focusing on *in vivo* studies are needed to strengthen a promising therapeutic role of TRPML1 agonists for JNCL.

It remains to be elucidated why besides main breakdown products of the glycerophospholipids including GPDs and lyso-glycerophospholipids^15^ lipids like Gb3 accumulate in CLN3 deficient cells. Lysosomal proteome profiling in a CLN3 mutant cerebellar granule cell line revealed decreased levels of several lysosomal phospholipases^58^. Why this occurs needs clarification. CLN3-deficiency may result in trafficking defects of lysosomal enzymes^8^. Furthermore, many of these enzymes rely on BMPs which levels are greatly reduced in CLN3-KO lysosomes. These enzymes perhaps undergo premature lysosomal degradation. Finally, it can also not be excluded that collateral damage inside the lysosome occurs due to the accumulation of GPDs or lyso-glycerophospholipids. Since different therapeutics such as antisense oligonucleotides (ASOs)^50^, miglustat (ongoing clinical trial by Theranexus), gemfibrozil^51^ and tamoxifen^14^ are proposed to treat CLN3 disease, it will be important to understand what different treatments can achieve in terms of normalizing lysosomal defects that occur at different levels.

## Material and Methods

### Reagents

For cellular treatments the following reagents were used: Torin 1 (Tocris, 4247), Mitomycin C (Merck (Roche), 10107409001), ML-SA1 (Bio-Techne, 4746), ML-SA5 (Enamine Ltd., 2418670-70-7), Bafilomycin A (Sigma, B1793), ML-SI3 (Enamine Ltd., EN300-314172), Ionomycin (Cell Signaling, 9995).

For GPD synthesis, the following starting materials were purchases from Avanti: 18:1 Lyso PG (858125) 18:1 Lyso PE (846725), 18:1 LysoPI and 18:1 Lyso PS (858143)

### Generation and cultivation of CLN3-KO ARPE-19 cells

CLN3-KO ARPE-19 cells were generated using the CRISPR/Cas9 system. CLN3 targeting crRNA with the sequence ATCCCACTGACGAGAACCCG was mixed with tracrRNA to form crRNA:tracrRNA guide complexes. ARPE-19 cells were transfected with crRNA:tracrRNA duplex and Cas9 nuclease (IDT, 1074182) by electroporation. Single-cell clones were expanded and mutations were evaluated by PCR, BioAnalyzer and next generation sequencing (NGS). The ARPE-19 CLN3-KO line carries 8 nucleotide deletions in exon 3 on both alleles, leading to a premature stop codon and a truncated protein missing the lysosomal targeting sequence (c.[377-384del+383-390del]). On protein level, these are leading to changes from amino acid 127 onwards and result in a frameshift and a premature stop codon at amino acid 156 (p.[Q127fsX156+E127fsX156]).

ARPE-19 cells were cultured in DMEM/F-12 medium (Gibco, 11320-074), supplemented with 2 mM L-Glutamine (Life Technologies, 25030-024) and 10 % heat-inactivated FBS (Gibco, 10500-064).

To induce a cell cycle arrest, ARPE-19 cells were treated with 0.3 mM Mitomycin C for 2 hours and washed with PBS before replating in the required well-format. Cells were kept in culture for 7 days thereafter.

### Generation of TMEM192-3xHA expressing ARPE-19 cells

TMEM192-3xHA plasmid generation and lentivirus production and packaging were purchased from VectorBuilder. Fhe vector contained, in addition to the TMEM192-3xHA gene of interest, a UBC promoter and puromycin cassette for selection. ARPE-19 WT and CLN3-KO cells were infected with lentivirus using an MOI of 5. To facilitate infection 10 µg/ml polybrene (VectorBuilder) was co-applied with the virus, followed by a centrifugation at 930xg for 1h at 37°C. 24h after infection, 4 µg/ml of puromycin dihydrochloride (Gibco, A1113803) was added for selection. 48h after infection, cells were split, expanded and cultured as described above. TMEM192-3xHA expression and lysosomal localization were confirmed by immunoblot and immunocytochemistry analysis using HA (Novus, NB600-362, 1:4000) and LAMP2 (DSHB, H4B4, 1:500) antibodies.

### Immunocytochemistry, co-localization and live cell imaging

The following primary antibodies were used: ATPase subunit C (Abcam, ab181243, 1:2000), LAMP1 (Santa Cruz, H4A3, 1:350), LC3 (MBL, PM036, 1:400), LAMP2 (DSHB, H4B4, 1:500), NPC1 (Abcam, ab134113, 1:1000), TFEB (Cell Signaling, 4240, 1:100), goat anti-mouse IgG (H+L), CF^TM^ 647 (Sigma, SAB4600182, 1:1000), goat anti-rabbit IgG, (H+L), CF^TM^ 568 (Sigma, SAB46000085, 1:1000). Cellular nuclei were stained with DAPI (4′,6-Diamidino-2-phenylindole dihydrochloride, 1:2000).

Cells were fixed in 4% Paraformaldehyde (PFA/PBS) for 20 min at RT, washed twice and then incubated with 3% (w/v) BSA, 0.1% (w/v) Saponin, 0.1% (w/v) Triton-X in PBS (permeabilization/blocking buffer) for 1h. Cells were incubated with primary antibodies for 1-2 h, followed by secondary antibodies for 1h, both diluated in permeabilization/blocking buffer.

For live-cell imaging with LysoTracker green (DND-26, Thermo Fisher, Scientific, L7526), cells were treated with 0.1 µM LysoTracker and 1:1000 Hoechst (33342, Trihydrochloride, Trihydrate, Thermo Fisher Scientific, H1399) for 5 min at 37°C. Cells were then washed three times with PBS and imaged directly afterwards.

Imaging was performed with the Opera Phenix high-content screening system (Perkin Elmer) equipped with a 4.5MPixel camera. The image analysis was performed using ACapella 5.1, in a Columbus 2.9 environment via texture thresholding.

Independently, nuclear detection was done by DAPI or Hoechst staining. After intensity-based thresholding, the detected objects were further filtered by size, roundness and contrast.

Starting from the detected nuclei, the cell boundary was determined. Cell boundary detection was based on the cytoplasmic background signal of the other aforementioned stainings. For co-localization analysis, the intersection of detected texture masks was calculated. Co-localized objects were then used for mean cytoplasm intensity calculations of the corresponding channels. Then, co-localized objects in the cytoplasm were counted and divided by the cell count. Furthermore, the total area covered by co-localized objects divided by the area covered by detected aggregates of one or the other staining was calculated individually.

In case of TFEB, no texture thresholding was done. Instead, a ratio of mean TFEB intensity in the nucleus divided by mean TFEB intensity in the cytoplasm was calculated.

All aggregation methods were performed on a cell, then field, then well-level.

Bar graphs were assembled using Graph Pad Prism and statistically analyzed using the student’s t-test or a two-way ANOVA. Details including post hoc tests are denoted in the respective figure legend.

### BMP analysis using mass spectrometry

For sample preparation, cells were vortexed for 15 sec and sonicated for 1 min in 1 ml MeOH: H2O 5mM EGTA (2:1). After sonication, samples were transferred in 10 ml borosilicate tubes containing 10 ng of internal standard (BMP 14:0/14:0). 2.5 ml methanol and 2 ml water were added for liquid/liquid extraction followed by the addition of 2.5 ml dichloromethane. In order to avoid emulsion, 80 μl of saturated sodium chloride were added. The homogenate was vortexed for 5 min at RT, then centrifuged at 2500xg for 10 min. The lower organic phase was evaporated under a stream of nitrogen at 37°C. The dried residue was reconstituted in 100 μl methanol with 10 mM ammonium formate. 10 μl of the dissolved reconstitute was used for liquid chromatography and tandem mass spectrometry analysis (LC-MS/MS). A similar extraction protocol was used for analysis of supernatants. The liquid/liquid extraction was performed with 2 ml water.

#### Analytical Method

Samples were directly injected into an ACE3 C18 column (2.1 x 50 mm; 1.7 μm) from AIT (Cormeilles-en-Paris, France) with a water/methanol gradient. Mobile phase A was composed of water with 10 mM ammonium formate, whereas mobile phase B contained methanol with 10 mM ammonium formate. Samples were run with 95% phase B for 2 min, 95 to 100% phase B for 0.5 min and 100% phase B for 9.5 min. Flow rates were set to 0.4 ml/min from 0 to 2 min and 12.5 to 15 min and changed to 0.2 ml/min from 2.5 to 12 min.

Bis(monoacylglycerol)phosphate species (BMPs) and internal standard (BMP 14:0/14:0) were analyzed on a Quantum Ultra triple quadrupole (Thermo Electron Corporation, San Jose, CA, USA). Positive electrospray was performed on a Thermo IonMax ESI probe. To increase the sensitivity and specificity of the analysis, multiple reaction monitoring was used with following MS/MS transitions: BMP 28:0 MH+, 667,5-285,3; BMP 32:0 MH+, 723,5-313,5; BMP 32:1 MH+, 721,5-313,5; BMP 32:2 MH+, 719,5-311,5; BMP 34:0 MH+, 751,5-341,6; BMP 34:1 MH+, 749,5-339,3; BMP 34:2 MH+, 747,5-339,3; BMP 36:0 MH+, 779,6-341,6; BMP 36:1 MH+, 777,6-339,3; BMP 36:2 MH+, 775,5-339,3; BMP 36:3 MH+, 773,5-339,3; BMP 36:4 MH+, 771,5-337,5; BMP 38:1 MH+, 805,6-339,3; BMP 38:2 MH+, 803,6-339,3; BMP 38:3 MH+, 801,6-339,3; BMP 38:4 MH+, 799,5-339,3; BMP 38:5 MH+, 797,5-339,3; BMP 38:6 MH+, 795,5-337,5; BMP 40:1 MH+, 833,6-339,3; BMP 40:2 MH+, 831,6-339,3; BMP 40:3 MH+, 829,6-339,3; BMP 40:4 MH+, 827,6-339,3; BMP 40:5 MH+, 825,6-339,3; BMP 40:6 MH+, 823,5-339,3; BMP 40:7 MH+, 821,5-339,3; BMP 40:8 MH+, 819,5-361,5; BMP 40:10 MH+, 815,5-385,6; BMP 42:10 MH+, 843,5-361,5; BMP 44:10 MH+, 871,5-387,6; BMP 44:1 MH+, 869,5-387,6; BMP 44:12 MH+, 867,5-385,6. The annotation MH+ is indicating a measurement in positive ion mode, where M corresponds to the mass of the BMP and H to hydrogen. Collision energy for each transition was 30 eV and the tube lens had 170 arbitrary units.

Following spray chamber settings were used: heated capillary, 400°C; spray voltage, 5000 V; sheath gas, 40 arbitrary units; auxiliary gas 10 arbitrary units. Calibration curves were produced by using synthetic BMP 18:1/18:1 and BMP 14:0/14:0 (Avanti Polar Lipid, Alabaster, AL, USA). The amounts of BMP 18:1/18:1 in the samples were determined by using inverse linear regression of standard curves. The relative quantification of the BMP species without analytical standard was performed using BMP 18:1/18:1 calibration curve.

### Lyso-IP and GPD analysis using mass spectrometry

#### LysoIP

Lysosome purification by LysoIP was adapted from the method previously described (Abu-Remaileh et al., 2017) and optimized for the ARPE-19 cell line. Briefly, cells were grown to 95% confluency in 15 cm cell culture dishes. For the cell cycle arrest and drug treatment, cells were grown to 80% confluency. If lysosomal isolates were to be used for downstream metabolomics analysis, 4 μM Lysotracker Red DND-99 (Invitrogen) was added to each plate 1 hour prior to cell harvest. At the time of harvest, each plate was washed 2x with 5 ml of ice-cold PBS and cells were scraped in 1 ml ice cold KPBS (136 mM KCl, 10 mM KH2PO4, pH 7.25 in Optima LC–MS water) and transferred to a 2 ml tube. From cell harvest on everything was performed on ice. Cells were pelleted by centrifugation at 1000*xg* for 2 min at 4 °C. Following centrifugation, cells were resuspended in 500 μl fresh KPBS and 12.5 μl of the cell suspension was taken for whole-cell fraction analysis. The remaining cell suspension was lysed by trituration using a 29.5-gauge insulin syringe (EXELINT international), then diluted with an additional 500 μl KPBS and centrifuged at 1000*xg* for 2 min at 4°C to pellet cell debris. The lysosome-containing supernatant (1ml) was added to tubes containing an equivalent of 100 μl Pierce anti-HA magnetic beads (Thermo Scientific) and rocked for 3 min at 4 °C. Following incubation, the bead-lysosome complexes were washed 3x with ice cold KPBS. For metabolite extraction, lysosome and whole-cell fractions were extracted in 50 μl and 225 μl, respectively, of 80% methanol in LC–MS water containing 500 nM isotope-labelled amino acids used as internal standards (Cambridge Isotope Laboratories). Following extraction, HA-binding beads were removed from the lysosome fractions using a rotary magnet and cell debris was removed from whole-cell extractions by centrifugation at 20,000*xg* at 4 °C for 15 min. The metabolomic extractions were analyzed on the Agilent Triple-Quadrupole 6470 Mass Spectrometer with workflow and parameters described below.

#### GPDs quantitation

Metabolites were separated on a SeQuant® ZIC®-pHILIC 50 x 2.1 mm column (Millipore Sigma 1504590001) with a 20 x 2.1 mm (Millipore Sigma 1504380001) guard and connected to a 1290 LC system to carry out hydrophilic interaction chromatography (HILIC) for metabolite separation prior to mass spectrometry. The LC system was coupled to the 6470A triple quadrupole (QQQ) mass spectrometer equipped with an LC-ESI probe. External mass calibration was performed using the standard calibration mixture every 7 days. Injection volumes of 1-2.5 µl were used for each sample, with single injection for both positive and negative ionization modes. Mobile phases: A, 20 mM ammonium carbonate and 0.1% ammonium hydroxide dissolved in 100% LC/MS grade water; B, 100% LC/MS grade acetonitrile. The chromatographic gradient was adapted from Laqtom et al. 2022. In brief, the elution was performed with a gradient of 10 min; linear decrease from 80-20% B from 0-7 min; fast linear increase from 20-80% B from 7-7.5 min; 80% B hold from 7.5-10 min. The flow rate was 0.15 ml/min. The column compressor and autosampler were held at 55 °C and 4 °C, respectively. The mass spectrometer parameters were as follows: the spray voltage was set to 3.5 kV in positive mode and 2.5 kV in negative mode, and the gas temperature and the sheath gas flow were held at 250 °C and 300 °C, respectively. Both gas flow and sheath gas flow were 12 L/min while the nebulizer was maintained at 25 psi. These conditions were held constant for both positive- and negative-ionization mode acquisitions.

The mass spectrometer was operated in Multiple Reaction Method (MRM) for targeted analysis of species of interest. Standard GPC was purchased from Cayman chemical and the four other GPDs (GPE, GPG, GPI and GPS) were chemically synthesized (Laqtom et al., 2022) from corresponding glycerophospholipids purchased from Avanti polar lipids. The GPDs and internal standards were optimized using a MassHunter Optimizer MRM. MassHunter Optimizer MRM is an automated method development software used to generate and optimize MRM transitions accumulating at most 4 products with different abundances from singly ionized species. The two most abundant transitions from either the negative or positive mode were selected to detect each species. The GPDs precursor-product ion pairs (m/z) used for MRM of the compounds were as follows: GPC: 258.1→104.1/86.1; GPE: 214.1→140.0/79.0; GPG: 247.1→172.9/99.0; GPI: 333.1→153.0/79.0 and GPS: 258→171.0/79.0.

High-throughput annotation and relative quantification of lipids were performed using a qualitative analysis software of MassHunter acquisition data and QQQ quantitative analysis (Quant-My-Way) software. Individual lipid species shown in the figures were validated using the Qualitative software by manually checking the peak alignment and matching the retention times and MS/MS spectra to the characteristic fragmentation compared to the standard compounds. Analyzing two transitions for the same compound and looking for similar relative response was an added validation criterion to ensure the correct species were identified and quantified. The raw abundances of the internal controls (iso-labeled amino acids) were checked as additional control as well as endogenous amino acids. The MRM method and retention time were used to quantify all lipid species using the quantification software, and the raw peak areas of all species were exported to Microsoft Excel for further analysis. The average of blanks was subtracted from all raw abundances and the sample divided by the average of the WT strain. For the drug treatment after cell cycle block, each sample was divided by the average of the WT strain treated with DMSO.

#### Electron microscopy

ARPE-19 WT and CLN3-KO cells were fixed with 1% wt/vol glutaraldehyde and 4% wt/vol paraformaldehyde in 0.1 M PB, pH 7.2 overnight. Thereafter the cells were spun down 3 times at 1000xg after each washing step with PBS. The pellets were embedded in 3% agarose (Invitrogen) and small pieces were cut and rinsed three times in 0.1 M sodium cacodylate buffer (pH 7.2–7.4). The cell pieces were incubated in 1% osmium tetroxide in cacodylate buffer for 20 minutes on ice. After osmication, the cells were dehydrated using ascending ethyl alcohol concentration steps, followed by two rinses in propylene oxide. Infiltration of the embedding medium was performed by immersing the samples in a 1:1 mixture of propylene oxide and Epon and finally in neat Epon and polymerized at 60 °C. Semithin sections (0.5 µm) from cells were prepared for light microscopy mounted on glass slides and stained for 1 minute with 1% Toluidine blue. Ultrathin sections (60nm) were examined in an EM902 (Zeiss, Germany). Pictures were taken with a MegaViewIII digital camera (A. Tröndle, Moorenweis, Germany).

#### Immunoblotting

The following primary antibodies were used: LAMP1 (Abcam, ab24170, 1:1000), NPC1 (Abcam, ab134113, 1:2000), LAMP2 (DSHB, H4B4, 1:500), LC3 (Cell Signaling, 2775, 1:500), TFEB (Cell Signaling, 4240, 1:500), Cathepsin B (R&D, AF953, 1:800), Cathepsin D (abcam, ab75852, 1:5000), TPP1 (Abcam, ab195234, 1:1000), Vinculin (Abcam, ab129002, 1:5000), GAPDH (Sigma, G9545, 1:10000). As secondary antibodies, horseradish peroxidase (HRP) conjugated antibodies from Dako (P0448 and P0447) were used with the dilution specified on the instruction leaflet.

Cells were dissociated in 0.05% Trypsin-EDTA (Thermo-Fisher, 25300-054), washed with PBS and lysed in RIPA buffer (Sigma, R0278) containing 1x protease and phosphatase inhibitor cocktail (Roche, cOmplete Tablets 04693159001, PhosStop 04906837001). Protein amounts were quantified using the BCA protein assay (Thermo Fisher Scientific, 23227). Between 20-30 µg protein was used per sample. Protein samples were mixed with LDS buffer (Novex, B0007), Sample Reducing Agent (Life technologies, NP0009) and filled up with RIPA buffer to the same volume. Denaturation was performed at 95°C for 5 min or 70°C at 10 min in case of membrane proteins. For LC3 protein, Tris-Glycine gels, for all other proteins, Bis-Tris gels from Thermo Fisher Scientific ranging from 4-16% were used. Proteins were immunoblotted on 0.2 µm nitrocellulose membranes (GE Healthcare Amersham, 10600006) using Pierce Methanol-free Transfer buffer (Thermo Fisher Scientific, 35040). Blocking, primary and secondary antibody incubations were done in TBS-T (Roth, 1061.1) supplemented with 5% milk powder (Heirler, 4010318030305) or 5% BSA (Sigma, A7906). Blots were blocked for 1h at RT and primary antibody was added over night at 4°C. After three washing steps with TBS-T, each at least for 10 min, blots were incubated for 1h with secondary antibodies followed by another 3 washing steps with TBS-T. Proteins were visualized via chemiluminescence using Pierce ECL Western blotting substrate (Thermo Fisher Scientific, 32209). Protein bands were quantified using Fiji ImageJ.

#### mRNA expression analysis

Cells were treated as indicated. Dissociation of cells and RNA isolation was performed according to manual instructions using RNeasy Mini Kit (Qiagen, 74004). Reverse transcription into cDNA was performed using the SuperScript III First Strand kit (Invitrogen, 18080-051). The mastermix for cDNA synthesis contained in addition to RNA Oligo (dt) primers, random hexamers, and dNTP mix. 800 ng RNA were used for reverse transcription and 400 ng cDNA for RT-qPCR experiments. For real-time quantitative PCR (RT-qPCR) QuantiTect SYBR Green PCR Kit (Qiagen, 204145) and the LightCycler 480 system II from Roche were used. *PPIB, HPRT* and *NONO* were tested as reference genes. *PPIB* was chosen for normalization, since it was showing the most stable expression in all tested conditions. *PPIB* is validated as a suitable reference gene in other studies^52–54^. The following primers were used: *CLN3* fw: 5’-GGGTTCTCGTCAGTGGGATT-3’, rev: 5’-TAGCGAAGACCACACCACAC-3’; Quantitect primers from Qiagen: Hs_TFEB_1_SG QuantiTect Primer Assay (QT00069951), Hs_LAMP1_1_SG QuantiTect Primer Assay (QT00070994), Hs_LAMP2_1_SG QuantiTect Primer Assay (QT00077063), Hs_NPC1_1_SG QuantiTect Primer Assay (QT00066465), Hs_CTSB_1_SG QuantiTect Primer Assay (QT00088641), Hs_CTSD_1_SG QuantiTect Primer Assay (QT00020391), Hs_TPP1_1_SG QuantiTect Primer Assay (QT00097363), Hs_MAP1LC3B_1_SG QuantiTect Primer Assay (QT00055069). Relative mRNA expression levels were calculated based on the 2^-ΔΔCT^ method, where the target gene CT values were normalized to the reference gene (*PPIB*) and the untreated control (DMSO treated WT cells, without CCA).

#### Lysosomal patch clamp analysis

ARPE-19 cell lines were treated with 1 µM Apilimod (Axon, 1369) overnight before performing whole-endolysosomal patch-clamp. Currents were recorded using the EPC-10 patch-clamp amplifier (HEKA, Lambrecht, Germany) and PatchMaster acquisition software (HEKA). Data were digitized at 40kHz and filtered at 2.8kHz. For application of the TRPML1 agonist (ML-SA1) and antagonist (ML-SI3), cytoplasmic solution was completely exchanged by cytoplasmic solution containing agonists. Cytoplasmic solution contained 140 mM K-MSA, 5mM KOH, 4 mM NaCl, 0.39 mM CaCl₂, 1 mM EGTA and 10 mM HEPES (pH was adjusted with KOH to 7.2). The luminal solution contained 140 mM Na-MSA, 5 mM K-MSA, 2 mM Ca-MSA, 1 mM CaCl₂, 10 mM HEPES and 10 mM MES (pH was adjusted with methanesulfonic acid to 4.6), 500 ms voltage ramps from -100 to +100 mV were applied every 5 s in all of the patch-clamp experiments. All statistical analyses were evaluated using Origin9 and GraphPadPrism9.0 software.

#### TFEB knockdown

TFEB siRNAs (D-009798-03, Dharmacon) and ON-TARGET plus Non-targeting Pool used as scrambled siRNAs (D-001810-10-20, Dharmacon) were transfected into ARPE-19 cells using Lipofectamine RNAiMax transfection reagent (13778075, Invitrogen). Three days after inducing cell cycle arrest, 1 µM siRNAs were transfected into ARPE-19 cells according to manufacturers’ protocol. After an incubation time of four days, cells were treated with TRPML1 agonists, which were incubated for another 2 days until cells were fixed and analyzed via immunblotting and immunofluorescence analysis.

#### Lysosomal exocytosis

Compound treatment was performed in MEM medium containing 10 mM HEPES. Medium was discarded, cells were washed once with PBS and incubated with Trypsin-EDTA for 5 min at 37 °C. Double the amount of cold medium was added and the cell suspension was centrifuged at 200xg for 5 min at 4 °C. The cell pellet was resuspended in 1% BSA/ PBS containing LAMP1 antibody (Santa Cruz, H4A3, 1:400) and incubated for 20 min on a rotating wheel in the cold room. After centrifugation (200xg, 5 min, 4 °C), the cell pellet was resuspended in 1% BSA/ PBS containing secondary fluorescence antibody (PE anti-mouse IgG1, BioLegend, 406608) and incubated for 1 h as described above. After an additional centrifugation step, cell pellet was resuspended in 100 µl ice cold PBS and 0.2 µL Zombie NIR viability dye (BioLegend, 77184). Cells were kept on ice until their flow cytometry analysis using FACS Canto II from BD-Biosciences.

#### Statistical analysis

Sample sizes and n-number are denoted in the respective figure legend. Significance levels for comparison of groups were determined using an unpaired two-tailed Student’s *t*-test, a one-way or two-way ANOVA, depending on the data set. The respective post hoc analysis type is also specified in the figure legend. Details on statistical analysis including F-values and degrees of freedom are denoted in the figure legends. Analysis were performed using GraphPad Prism 8. Quantifications are plotted as means ± SD, if not otherwise stated.

## Data availability

The authors declare that all primary data are available within the article and its supplementary information files. All other data that support the findings of this study are available from the corresponding authors upon reasonable request.

## Acknowledgements

D.W. was supported by the NCL foundation in Hamburg. M.A.R., C.G., and E.K. have also received support from the NCL foundation. G.H. was supported by the Deutsche Forschungsgemeinschaft grant HE 3220/4-1. K.N. was supported by the Sarafan ChEM-H Chemistry/Biology Interface Program as a Kolluri Fellow and the Stanford Interdisciplinary Graduate Fellowship affiliated with the Wu Tsai Neurosciences Institute (Bio-X SIGF: Mark and Mary Steven’s Interdisciplinary Graduate Fellow). We thank the Metabolomics Knowledge Center (MKC) at Sarafan ChEM-H. A special thank goes to Herman van der Putten, research head of the NCL foundation, for enabling collaborations, scientific advice, many helpful discussions and help preparing the manuscript.

## Author contributions

Design and execution of experiments, data analysis and interpretation: D.W., R.T., K.N., N.N.L., M.S., C.W., S.M., D.G.S. and G.H. The ARPE-19 cell line was provided by C.G. Conceptualization, interpretation of data and editing the manuscript: D.W., S.M., D.G.S., G.H., M.A.R., C.G. and R.K. Design of the study and writing the manuscript D.W., D.G.S., and S.M. The final version of the manuscript was read and approved by all authors.

## References

1. Cotman, S. L. & Staropoli, J. F. The juvenile Batten disease protein, CLN3, and its role in regulating anterograde and retrograde post-Golgi trafficking. Clin. Lipidol. 7, 79–91 (2012).

2. Lerner, T. J. et al. Isolation of a novel gene underlying batten disease, CLN3. Cell 82, 949–957 (1995).

3. Cárcel-Trullols, J., Kovács, A. D. & Pearce, D. A. Cell biology of the NCL proteins: What they do and don’t do. Biochim. Biophys. Acta - Mol. Basis Dis. 1852, 2242–2255 (2015).

4. Mirza, M. et al. The CLN3 gene and protein: What we know. Mol. Genet. Genomic Med. 7, 1–41 (2019).

5. Luiro, K., Yliannala, K., Ahtiainen, L., Maunu, H. & Ja, I. Interconnections of CLN3, Hook1 and Rab proteins link Batten disease to defects in the endocytic pathway. Hum. Mol. Genet. 13, 3017–3027 (2004).

6. Vidal-Donet, J. M., Cárcel-Trullols, J., Casanova, B., Aguado, C. & Knecht, E. Alterations in ROS Activity and Lysosomal pH Account for Distinct Patterns of Macroautophagy in LINCL and JNCL Fibroblasts. PLoS One 8, (2013).

7. Uusi-Rauva, K. et al. Neuronal ceroid lipofuscinosis protein CLN3 interacts with motor proteins and modifies location of late endosomal compartments. Cell. Mol. Life Sci. 69, 2075–2089 (2012).

8. Yasa, S. et al. CLN3 regulates endosomal function by modulating Rab7A-effector interactions. J. Cell Sci. 133, 1–14 (2020).

9. Klein, M. et al. Converging roles of PSENEN/PEN2 and CLN3 in the autophagy-lysosome system. Autophagy 0, 1–18 (2021).

10. Oetjen, S., Kuhl, D. & Hermey, G. Revisiting the neuronal localization and trafficking of CLN3 in juvenile neuronal ceroid lipofuscinosis. J. Neurochem. 139, 456–470 (2016).

11. Nugent, T., Mole, S. E. & Jones, D. T. The transmembrane topology of Batten disease protein CLN3 determined by consensus computational prediction constrained by experimental data. FEBS Lett. 582, 1019–1024 (2008).

12. Ratajczak, E., Petcherski, A., Ramos-Moreno, J. & Ruonala, M. O. FRET-assisted determination of CLN3 membrane topology. PLoS One 9, 21–23 (2014).

13. Fossale, E. et al. Membrane trafficking and mitochondrial abnormalities precede subunit c deposition in a cerebellar cell model of juvenile neuronal ceroid lipofuscinosis. BMC Neurosci. 5, 1–13 (2004).

14. Soldati, C. et al. Repurposing of tamoxifen ameliorates CLN 3 and CLN 7 disease phenotype. EMBO Mol. Med. e13742, 1–19 (2021).

15. Laqtom, N. N. et al. CLN3 is required for the clearance of glycerophosphodiesters from lysosomes Check for updates. Nature (2022).

16. Hobert, J. A. & Dawson, G. A novel role of the Batten disease gene CLN3: Association with BMP synthesis. Biochem. Biophys. Res. Commun. 358, 111–116 (2007).

17. Tang, C. et al. A human model of Batten disease shows role of CLN3 in phagocytosis at the photoreceptor–RPE interface. Commun. Biol. 4, 1–18 (2021).

18. Zhong, Y. et al. Loss of CLN3, the gene mutated in juvenile neuronal ceroid lipofuscinosis, leads to metabolic impairment and autophagy induction in retina pigment epithelium. Biochim. Biophys. Acta - Mol. Basis Dis. 1866, 165883 (2020).

19. Tedeschi, V. et al. The activation of Mucolipin TRP channel 1 (TRPML1) protects motor neurons from L-BMAA neurotoxicity by promoting autophagic clearance. Sci. Rep. 9, 1–11 (2019).

20. Cao, Q. et al. BK Channels Alleviate Lysosomal Storage Diseases by Providing Positive Feedback Regulation of Lysosomal Ca2+ Release. Dev. Cell 33, 427–441 (2015).

21. Wang, W. et al. Up-regulation of lysosomal TRPML1 channels is essential for lysosomal adaptation to nutrient starvation. Proc. Natl. Acad. Sci. U. S. A. 112, E1373–E1381 (2015).

22. Yu, L. et al. Small-molecule activation of lysosomal TRP channels ameliorates Duchenne muscular dystrophy in mouse models. Sci. Adv. 6, (2020).

23. Shen, D. et al. Lipid storage disorders block lysosomal trafficking by inhibiting a TRP channel and lysosomal calcium release. Nat. Commun. 3, (2012).

24. Bae, M. et al. Activation of TRPML1 clears intraneuronal Aβ in preclinical models of HIV infection. J. Neurosci. 34, 11485–11503 (2014).

25. Cao, Q., Yang, Y., Zhong, X. Z. & Dong, X. P. The lysosomal Ca2+ release channel TRPML1 regulates lysosome size by activating calmodulin. J. Biol. Chem. 292, 8424–8435 (2017).

26. Li, X. et al. A molecular mechanism to regulate lysosome motility for lysosome positioning and tubulation. Nat. Cell Biol. 18, 404–417 (2016).

27. Scotto Rosato, A., et al. TRPML1 links lysosomal calcium to autophagosome biogenesis through the activation of the CaMKKβ/VPS34 pathway. Nat. Commun. 10, (2019).

28. Medina, D. L. et al. Lysosomal calcium signalling regulates autophagy through calcineurin and TFEB. Nat. Cell Biol. 17, 288–299 (2015).

29. Lojewski, X. et al. Human iPSC models of neuronal ceroid lipofuscinosis capture distinct effects of TPP1 and CLN3 mutations on the endocytic pathway. Hum. Mol. Genet. 23, 2005–2022 (2014).

30. Scotto Rosato, A., et al. TPC 2 rescues lysosomal storage in mucolipidosis type IV, Niemann – Pick type C 1, and Batten disease. EMBO Mol. Med. e15377, (2022).

31. Hongo, K. et al. Massive accumulation of globotriaosylceramide in various tissues from a Fabry patient with a high antibody titer against alpha-galactosidase A after 6 years of enzyme replacement therapy. Mol. Genet. Metab. Reports 24, 100623 (2020).

32. Kobayashi, T. et al. A lipid associated with the antiphospholipid syndrome regulates endosome structure and function. Nature 392, 193–197 (1998).

33. Brotherus, J. & Renkonen, O. Subcellular distributions of lipids in cultured BHK cells: evidence for the enrichment of lysobisphosphatidic acid and neutral lipids in lysosomes. J. Lipid Res. 18, 191–202 (1977).

34. Gallala, H. D. & Sandhoff, K. Biological function of the cellular lipid BMP-BMP as a key activator for cholesterol sorting and membrane digestion. Neurochem. Res. 36, 1594–1600 (2011).

35. Gruenberg, J. Life in the lumen: The multivesicular endosome. Traffic 21, 76–93 (2020).

36. Boland, S. et al. Deficiency of the frontotemporal dementia gene GRN results in gangliosidosis. Nat. Commun. 13, (2022).

37. Ilnytska, O. et al. Enrichment of NPC1-deficient cells with the lipid LBPA stimulates autophagy, improves lysosomal function, and reduces cholesterol storage. J. Biol. Chem. 297, 100813 (2021).

38. Moreau, D. et al. Drug-induced increase in lysobisphosphatidic acid reduces the cholesterol overload in Niemann–Pick type C cells and mice. EMBO Rep. 20, 1– 15 (2019).

39. Wang, W. et al. Up-regulation of lysosomal TRPML1 channels is essential for lysosomal adaptation to nutrient starvation. Proc. Natl. Acad. Sci. U. S. A. 112, E1373–E1381 (2015).

40. Samie, M. et al. A TRP Channel in the Lysosome Regulates Large Particle Phagocytosis via Focal Exocytosis. Dev. Cell 26, 511–524 (2014).

41. Palmieri, M. et al. Characterization of the CLEAR network reveals an integrated control of cellular clearance pathways. Hum. Mol. Genet. 20, 3852–3866 (2011).

42. Cao, Y. et al. Autophagy is disrupted in a knock-in mouse model of juvenile neuronal ceroid lipofuscinosis. J. Biol. Chem. 281, 20483–20493 (2006).

43. Settembre, C. et al. TFEB Links Autophagy to Lysosomal Biogenesis. Science (80-.). 332, 1429–1433 (2011).

44. Klionsky, D. J. et al. Guidelines for the use and interpretation of assays for monitoring autophagy (3rd edition). Autophagy 12, 1–222 (2016).

45. Medina, D. L. et al. Transcriptional activation of lysosomal exocytosis promotes cellular clearance. Dev. Cell 21, 421–430 (2011).

46. Andrews, N. W. Detection of Lysosomal Exocytosis by Surface Exposure of Lamp1 luminal Epitopes. Lysosomes Methods Protoc. Mehtods Mol. Biol. 1594, 366–419 (2017).

47. Mole, S. E. et al. Clinical challenges and future therapeutic approaches for neuronal ceroid lipofuscinosis. Lancet Neurol. 18, 107–116 (2019).

48. Wright, G. A. et al. Juvenile Batten Disease (CLN3): Detailed Ocular Phenotype, Novel Observations, Delayed Diagnosis, Masquerades, and Prospects for Therapy. Ophthalmol. Retin. 4, 433–445 (2020).

49. Dang Do, A. N., et al. Neurofilament light chain levels correlate with clinical measures in CLN3 disease. Genet. Med. 23, 751–757 (2021).

50. Centa, J. L. et al. Therapeutic efficacy of antisense oligonucleotides in mouse models of CLN3 Batten disease.

51. J. Malabendu, D. Debashis, P. Jit, P. K. Activation of PPARα exhibits therapeutic efficacy in a mouse model of juvenile neuronal ceroid lipofuscinosis. J. Neurosci. JN-RM-2447–21 (2023) doi:10.1523/JNEUROSCI.2447-21.2023.

52. Jin, E. et al. Evidence of a novel gene HERPUD1 in polypoidal choroidal vasculopathy. Int. J. Clin. Exp. Pathol. 8, 13928–13944 (2015).

53. Giri, A. & Sundar, I. K. Evaluation of stable reference genes for qPCR normalization in circadian studies related to lung inflammation and injury in mouse model. Sci. Rep. 12, 1–14 (2022).

54. Pachot, A., Blond, J. L., Mougin, B. & Miossec, P. Peptidylpropyl isomerase B (PPIB): A suitable reference gene for mRNA quantification in peripheral whole blood. J. Biotechnol. 114, 121–124 (2004).

