## Supplementary Material for "TRPML1 activation ameliorates lysosomal phenotypes in CLN3 deficient retinal pigment epithelial cells"

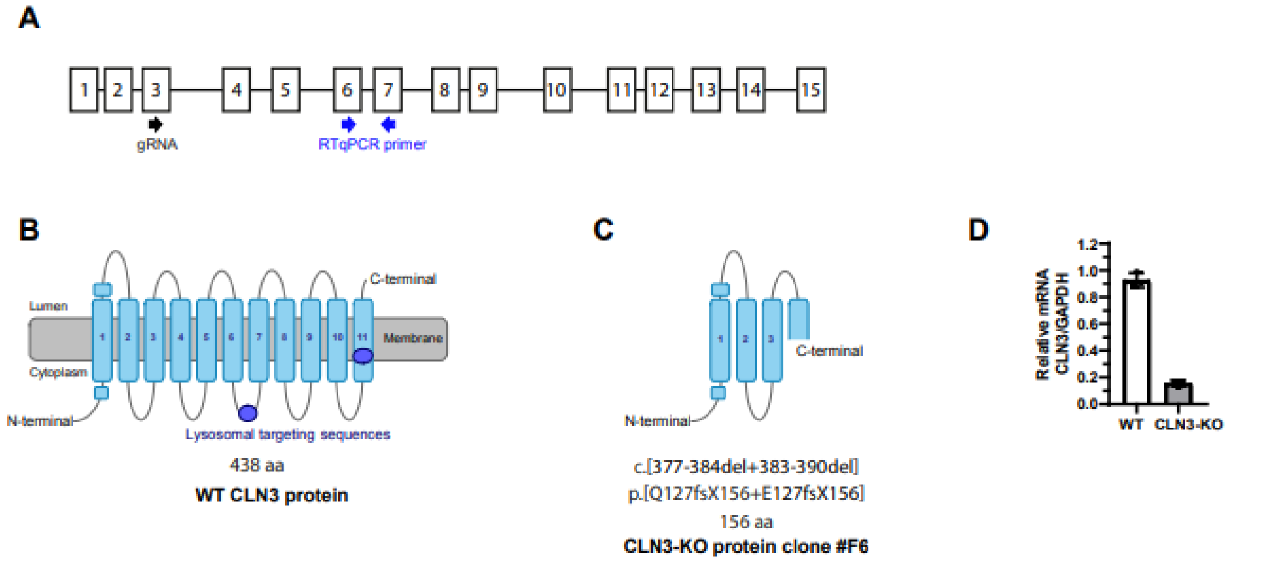


**Figure S1: ARPE-19 CLN3-KO model. A** CLN3 gene scheme, boxes represent exons, the position of guide RNA (gRNA) used to generate ARPE-19 CLN3-KO cell line via CRISPR gene editing is indicated. qPCR primers to determine knockout transcript levels are depicted. **B, C** Predicted CLN3 protein topology with 11 transmembrane structure published on alphafold: WT protein (**B**) and the truncated protein encoded in CLN3-KO ARPE-19 cells (**C**). The CLN3-KO line carries several nucleotide deletions on cDNA level (c.[377-384del+383-390del]), leading to a frameshift and a premature stop codon at amino acid 156 (p.[Q127fsX156+E127fsX156]). **D** Relative mRNA expression of CLN3 without cell cycle arrest, normalized to GAPDH. The plot summarizes three biological replicates with each three technical replicates from one experiment.

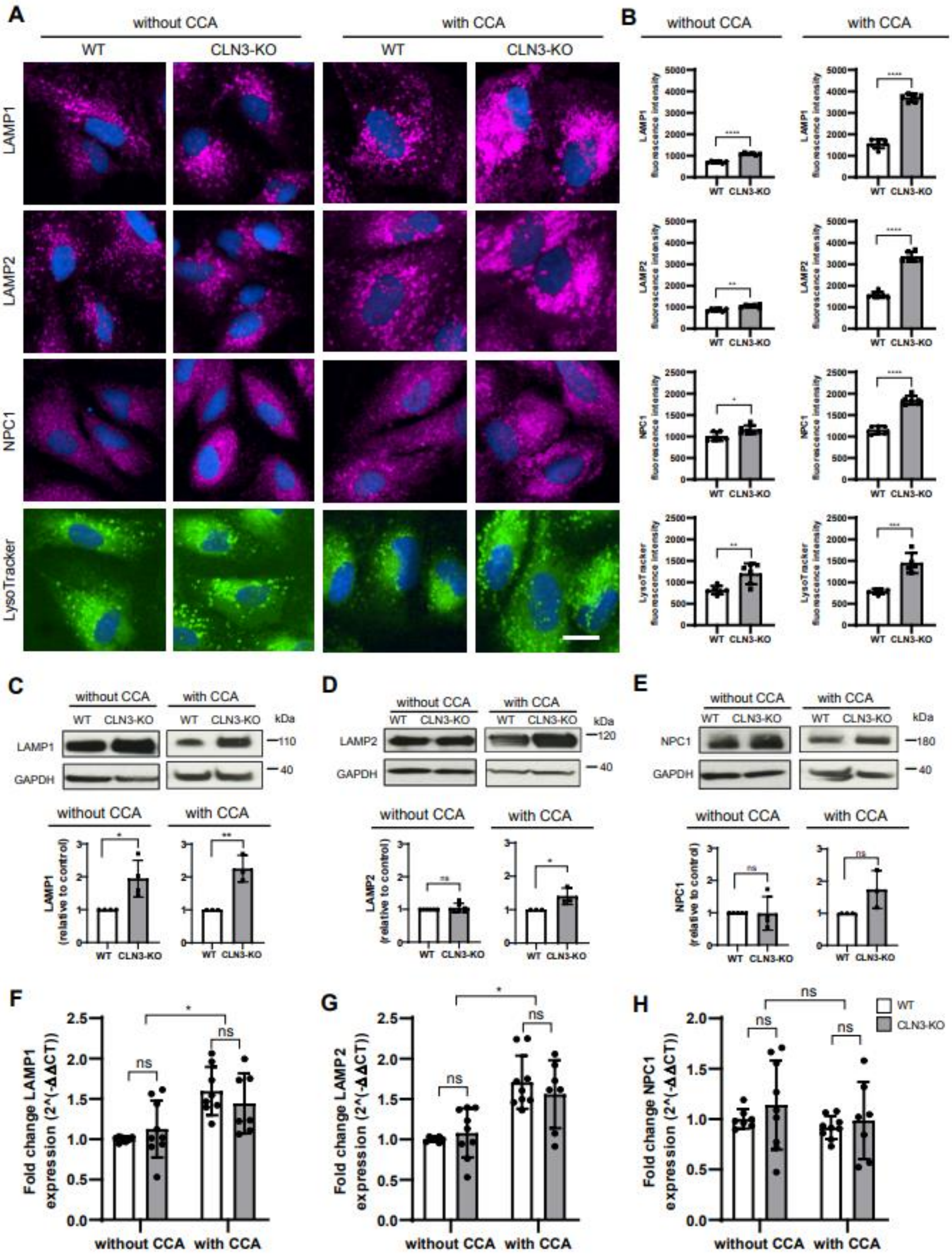


**Figure S2: Lysosomal marker analysis on protein and gene expression level**. **A** Representative immunocytochemical images of LAMP1, LAMP2, NPC1 and LysoTracker in ARPE-19 WT and CLN3-KO cells with and without cell cycle arrest (CCA). Scale bar: 20 µm. **B** Quantification of fluorescence intensities from one experiment (images shown in A). In total three independent experiments were performed. Values are means ± SD. P-values calculated by unpaired two-tailed Student's *t*-test **C, D, E** Representative images of immunoblot analysis of LAMP1 (**C**), LAMP2 (**D**) and NPC1 (**E**) with and without CCA. Plots show quantification of LAMP1, LAMP2 and NPC1 protein levels, normalized to GAPDH. Values are means ± SD, *n*=3 independent experiments. P-values calculated by unpaired two-tailed Student's *t*-test. **F, G, H** Relative mRNA expression levels of LAMP1 (**F**), LAMP2 (**G**) and NPC1 (**H**), normalized to Peptidyl-prolyl cis-trans isomerase B (PPIB) WT cells without CCA. With and without cell cycle arrest. Values are means ± SD, n=3 independent experiments. P-values were calculated with a two-way ANOVA coupled with multiple comparisons using Sidak test. ns, nonsignificant<0.1234, **p*-value<0.0332; ***p*<0.0021, ****p*-value<0.0002, *****p*-value<0.0001.

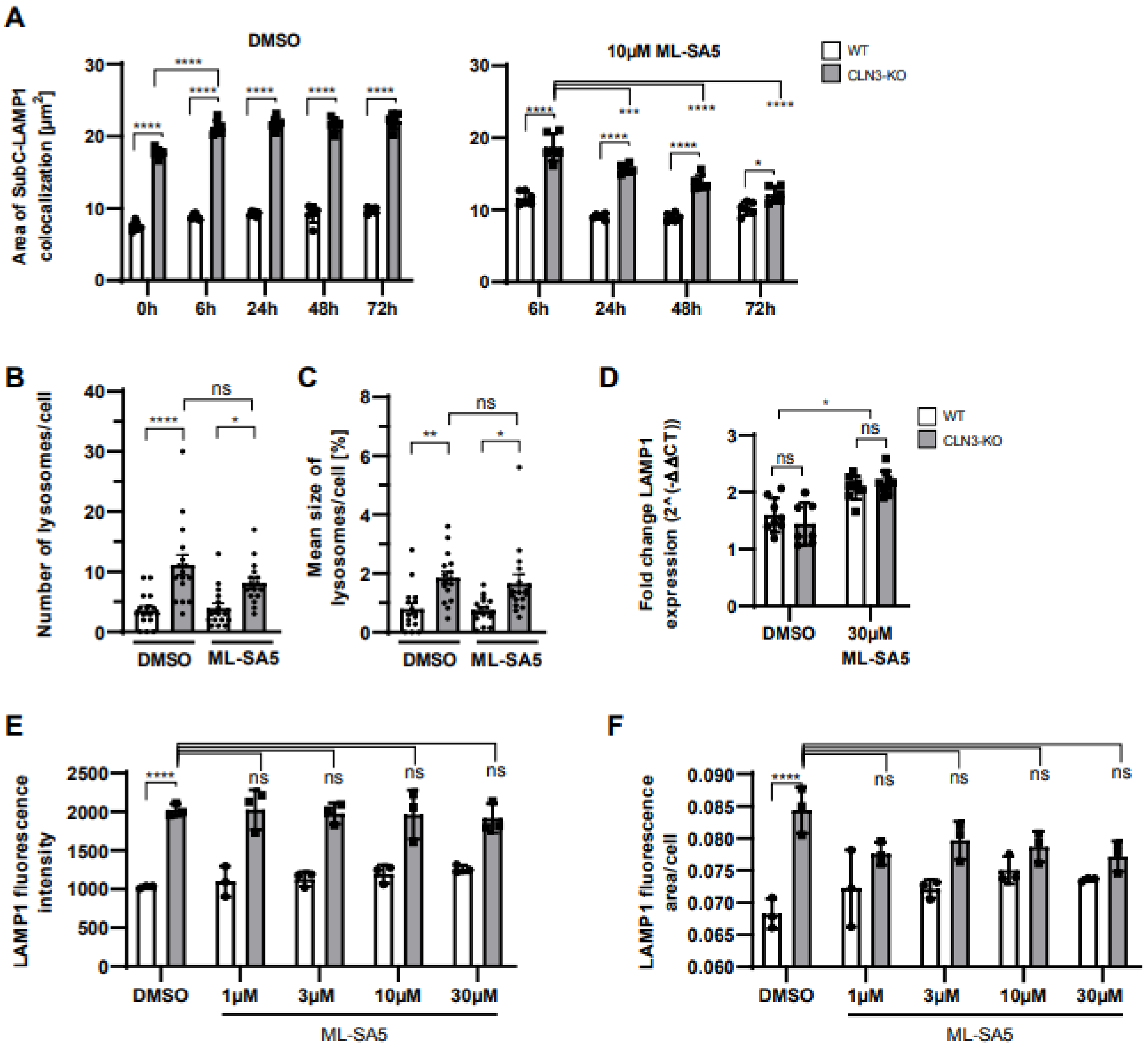


**Figure S3: Effects of TRPML1 activation on lysosomes, LAMP1 and SubC levels. A** Plots show area of SubC-LAMP1 colocalization, supplementary data to Fig. 4 A-C. Cell cycle arrested cells were treated with 0.3 % DMSO (vehicle) and 10 µM ML-SA5 for different time periods. Summarized data from one experiment. In total three independent experiments were performed. Values are means ± SD. P-values were calculated by two-way ANOVA coupled with multiple comparisons using Tukey test. **B, C** Plots show quantification of lysosomal number (**B**) and size (**C**) summarized from 16 electron microscopy images. Cell cycle arrested cells were treated with 10 µM ML-SA5 for 48 h. Values are means ± SEM. P-values were calculated by a one-way ANOVA coupled with multiple comparisons using

Tukey test. **D** Relative mRNA expression levels of LAMP1 normalized to PPIB and WT DMSO treated cells without CCA. Cells were treated for 24 h with 30 µM ML-SA5, with cell cycle arrest. Values are means ± SD, n=3 independent experiments. P-values were calculated by a two-way ANOVA coupled with multiple comparisons using Tukey test. **E, F** Plots show LAMP1 fluorescence intensity (**E**) and LAMP1 fluorescence area/cell (**F**) of cell cycle arrested cells treated for 48 h with DMSO/ML-SA5. Supplementary data to Fig. 4 A-C. Summarized data from one experiment. In total three independent experiments were performed. Values are means ± SD. P-values were calculated by a two-way ANOVA coupled with multiple comparisons using Tukey test. ns, nonsignificant<0.1234, **p*-value<0.0332; ***p*<0.0021, ****p*-value<0.0002, *****p*-value<0.0001.

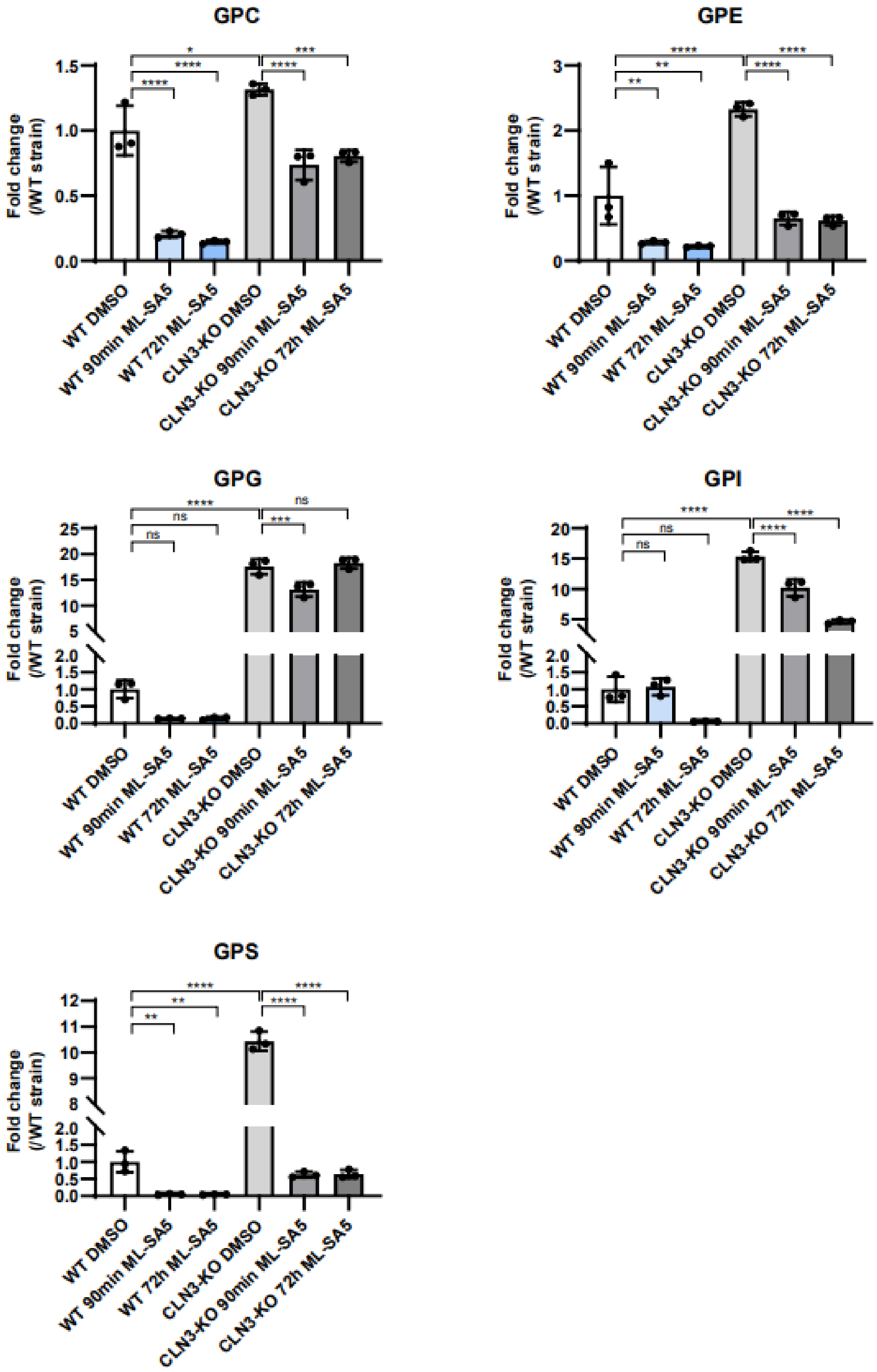


**Figure S4: Effect of TRPML1 activation on whole cell GPD levels in WT and CLN3-KO cells.** Plots show the fold change of the different GPD species from ARPE-19 WT and CLN3-KO whole cells under CCA and treatment with 10 µM ML-SA5 for 90min and 72h. Values are means ± SD of three independent experiments. P-values were calculated by one-way ANOVA coupled with multiple comparisons using Tukey test. ns, nonsignificant<0.1234, *$p$-value<0.0332; **$p<0.0021$, ***$p$-value<0.0002, ****$p$-value<0.0001.

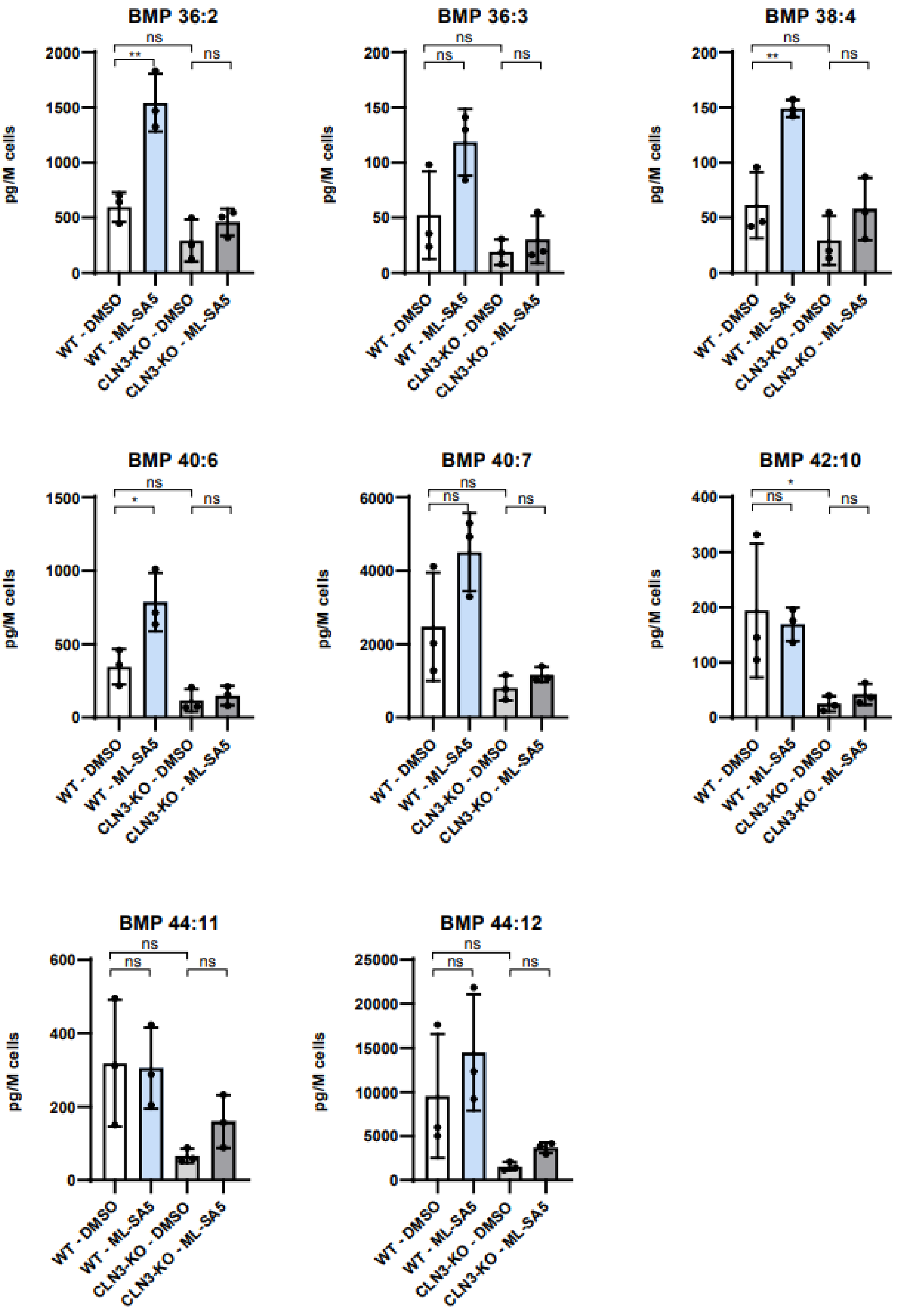

BMP 36:2
BMP 36:3
BMP 38:4
BMP 40:6
BMP 40:7
BMP 42:10
BMP 44:11
BMP 44:12
pg/M cells
WT - DMSO
WT - ML-SA5
CLN3-KO - DMSO
CLN3-KO - ML-SA5
ns
**
*

**Figure S5: Effect of TRPML1 activation on BMP levels in non cell cycle arrested WT and CLN3-KO cells.** Plots show the abundance of BMP species in cell lysates of ARPE-19 WT and CLN3-KO cells with CCA and treatment with 3 µM ML-SA5 for 48h. Values are means ± SD of three independent experiments. P-values were calculated by one-way ANOVA coupled with multiple comparisons using Tukey test. ns, nonsignificant<0.1234, **p*-value<0.0332; ***p*<0.0021, ****p*-value<0.0002, *****p*-value<0.0001.

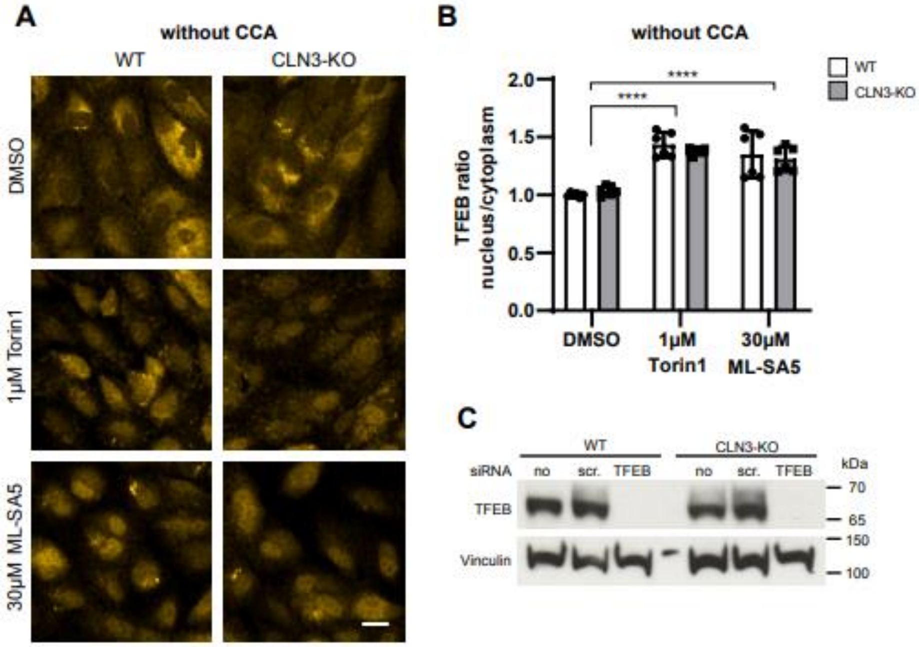


**Figure S6: Effect of TRPML1 activation on TFEB nuclear translocation in WT and CLN3-KO cells. A** Representative immunocytochemical images showing TFEB nuclear translocation after treatment of ARPE-19 WT and CLN3-KO cells without cycle arrest for 2 h with 1 µM Torin1 and 30 µM ML-SA5. **B** Quantification of images shown in A. Plot shows TFEB nuclear/cytosol ratio of one representative experiment. In total three independent experiments were performed. Values are means ± SD. **C** Representative immunoblot analysis of TFEB in cell cycle arrested ARPE-19 WT and CLN3-KO treated with no, scrambled (scr.) and TFEB siRNAs. *P*-values calculated by two-way ANOVA coupled with multiple comparisons using Sidak test. ns, nonsignificant<0.1234, **p*-value<0.0332; ***p*<0.0021, ****p*-value<0.0002, *****p*-value<0.0001. Scale bar: 20µm.

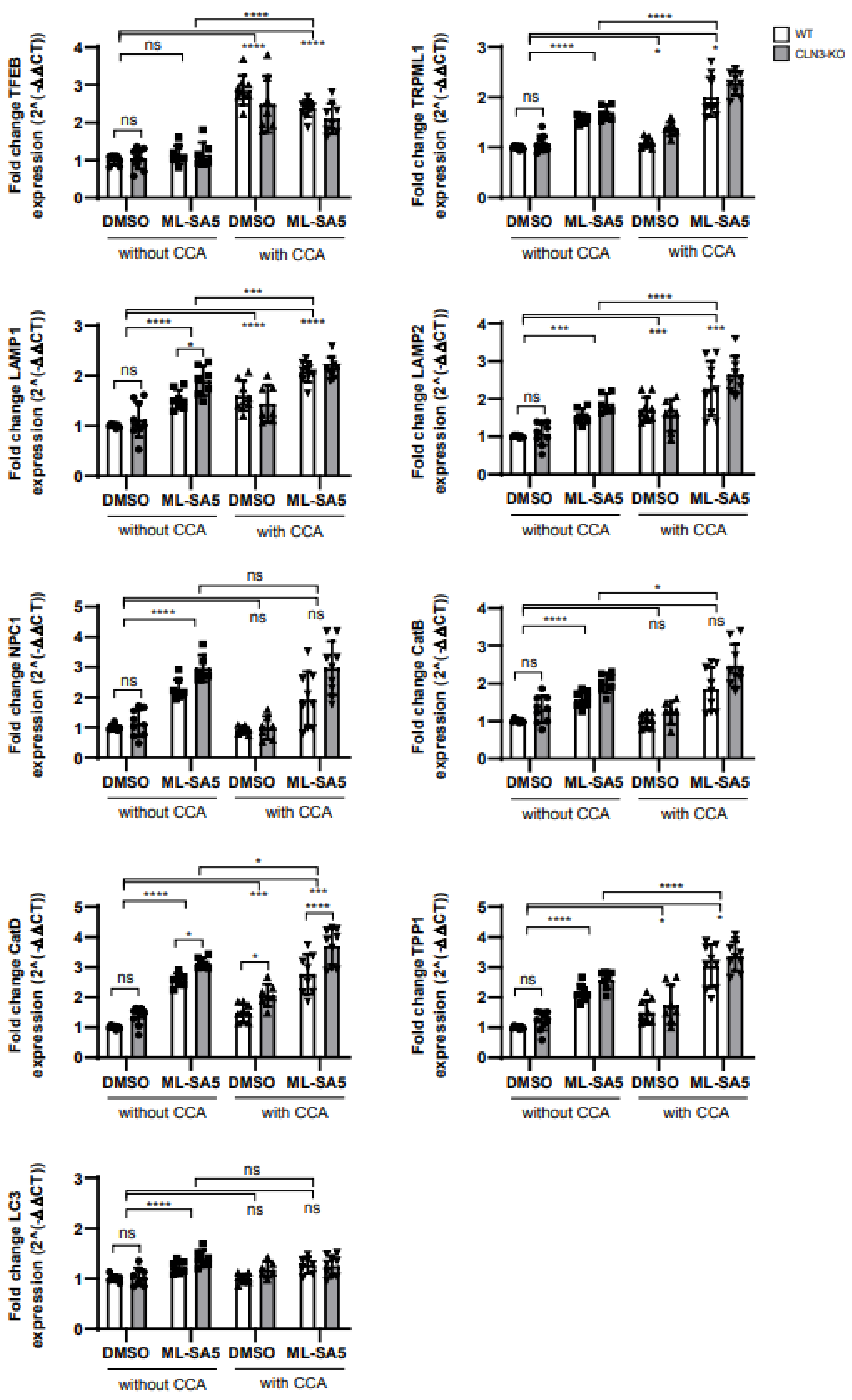
WT
CLN3-KO
Fold change TFEB expression (2^(-ΔΔCT))
Fold change TRPML1 expression (2^(-ΔΔCT))
Fold change LAMP1 expression (2^(-ΔΔCT))
Fold change LAMP2 expression (2^(-ΔΔCT))
Fold change NPC1 expression (2^(-ΔΔCT))
Fold change CatB expression (2^(-ΔΔCT))
Fold change CatD expression (2^(-ΔΔCT))
Fold change TPP1 expression (2^(-ΔΔCT))
Fold change LC3 expression (2^(-ΔΔCT))
DMSO
ML-SA5
without CCA
with CCA
ns
****
***
*

**Figure S7: Effect of TRPML1 activation on TFEB regulated genes.** Relative mRNA expression levels of TFEB, TRPML1, LAMP1, LAMP2, NPC1, CatB, CatD, TPP1 and LC3, all normalized to Peptidyl-prolyl cis-trans isomerase B (PPIB) and DMSO treated WT cells without CCA. With and without cell cycle arrest, with and without treatment of 30 µM ML-SA5 for 24h. Values are means ± SD, n=3 independent experiments. P-values were calculated with a two-way ANOVA coupled with multiple comparisons using Sidak test. ns, nonsignificant<0.1234, **p*-value<0.0332; ***p*<0.0021, ****p*-value<0.0002, *****p*-value<0.0001.

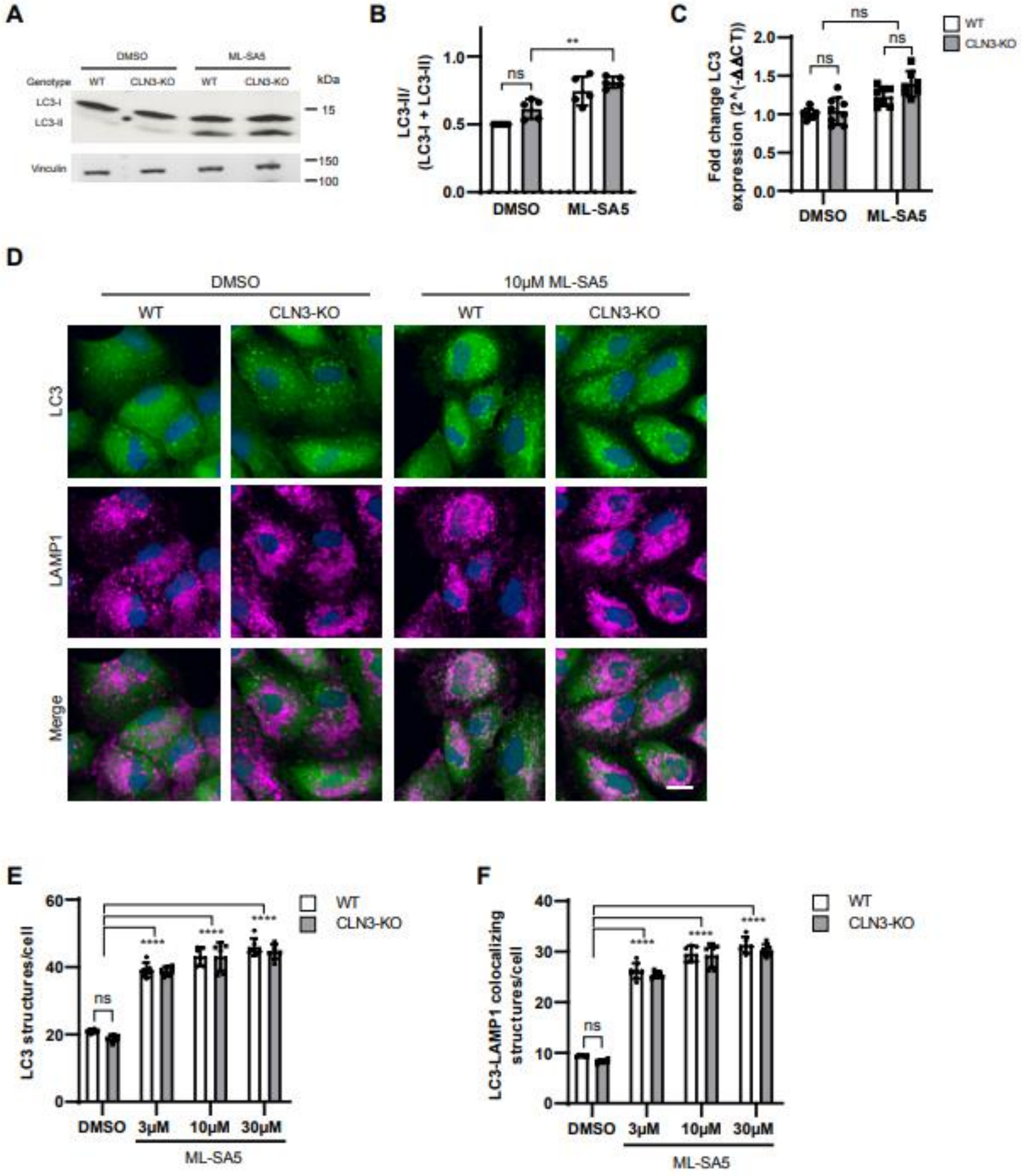

**Figure S8: Effect of TRPML1 activation on autophagy in non cell cycle arrested WT and CLN3-KO cells. A** Representative image of immunoblot analysis of LC3. Cells without cell cycle arrest were treated for 30 min with 30 µM ML-SA5. **B** Quantification of LC3-II/(LC3-I + LC3-II) levels normalized to Vinculin. Plot summarizes data from 5 independent experiments. **C** Relative mRNA expression of LC3, normalized to Peptidyl-prolyl cis-trans isomerase B (PPIB) and DMSO treated WT cells without CCA. Cells were treated for 24 h with 30 µM ML-SA5. Values are means ± SD, *n*=3 independent experiments. **D** Representative immunocytochemical images of LC3 and LAMP1 localization in ARPE-19 WT and CLN3-KO cells without cell cycle arrest. Scale bar: 20 µm. **E, F** Plots represent the number of LC3 structures/cell (**E**) and LC3-LAMP1 colocalizing structures/per cell (**F**). Cells were treated with for 30 min (**D-F**). Plots summarize data from one representative experiment. In total three independent experiments were performed. Values are means ± SD. *P*-values calculated by two-way ANOVA coupled with multiple comparisons using Tukey test. ns, nonsignificant<0.1234, **p*-value<0.0332; ***p*<0.0021, ****p*-value<0.0002, *****p*-value<0.0001.
